# A Splice Site-Sensing Conformational Switch in U2AF2 is Modulated by U2AF1 and its Recurrent Myelodysplasia-Associated Mutation

**DOI:** 10.1101/2019.12.31.891358

**Authors:** Chandani Warnasooriya, Eliezra Glasser, Callen F. Feeney, Kholiswa M. Laird, Jermaine L. Jenkins, Dmitri N. Ermolenko, Clara L. Kielkopf

## Abstract

An essential heterodimer of the U2AF1 and U2AF2 pre-mRNA splicing factors nucleates spliceosome assembly at polypyrimidine (Py) signals preceding the major class of 3ʹ splice sites. Among myelodysplastic syndromes (MDS), U2AF1 frequently acquires an S34F-encoding mutation. The influence of the U2AF1 subunit and its S34F mutation on the U2AF2 conformations remains unknown. Here, we employ single molecule Förster resonance energy transfer (FRET) to determine the influence of wild-type or S34F-substituted U2AF1 on the conformational dynamics of U2AF2 and its splice site RNA complexes. In the absence of RNA, the U2AF1 subunit stabilizes a high FRET value, which by structure-guided mutagenesis corresponds to a closed conformation of the tandem U2AF2 RNA recognition motifs (RRMs). When the U2AF heterodimer is bound to a strong, uridine-rich splice site, U2AF2 switches to a lower FRET value characteristic of an open, side-by-side arrangement of the RRMs. Remarkably, the U2AF heterodimer binds weak, uridine-poor Py tracts as a mixture of closed and open U2AF2 conformations, which are modulated by the S34F mutation. Shifts between open and closed U2AF2 may underlie U2AF1-dependent splicing of degenerate Py tracts and contribute to a subset of S34F-dysregulated splicing events in MDS patients.

## INTRODUCTION

During eukaryotic gene expression, introns are removed from the pre-mRNA transcripts to prepare the mature mRNA for translation. A multicomponent spliceosome machinery of proteins and small nuclear (sn)RNAs assembles in a tightly-controlled, stepwise process on the consensus sequences of pre-mRNA splice sites (1). At the major class of 3ʹ splice sites, a heterodimer of U2AF1 and U2AF2 (also called U2AF^35^ and U2AF^65^) recognizes an consensus AG-dinucleotide at the spliced junction and a preceding polypyrimidine (Py) tract (Figure 1A). This ribonucleoprotein complex recruits the U2 small nuclear ribonucleoprotein particle (snRNP) of the spliceosome. Whereas U2AF1 has a specific role in splicing so-called “AG”-dependent splice sites that lack a clear Py tract signal (2-4), the U2AF2 subunit is thought to serve general functions for splicing of uridine-rich sites (5). These different U2AF1-dependencies and degeneracy of the Py tract signal are thought to regulate alternative splicing of multi-exon genes, which generates numerous transcript variants in eukaryotes (6).

**Figure 1.**
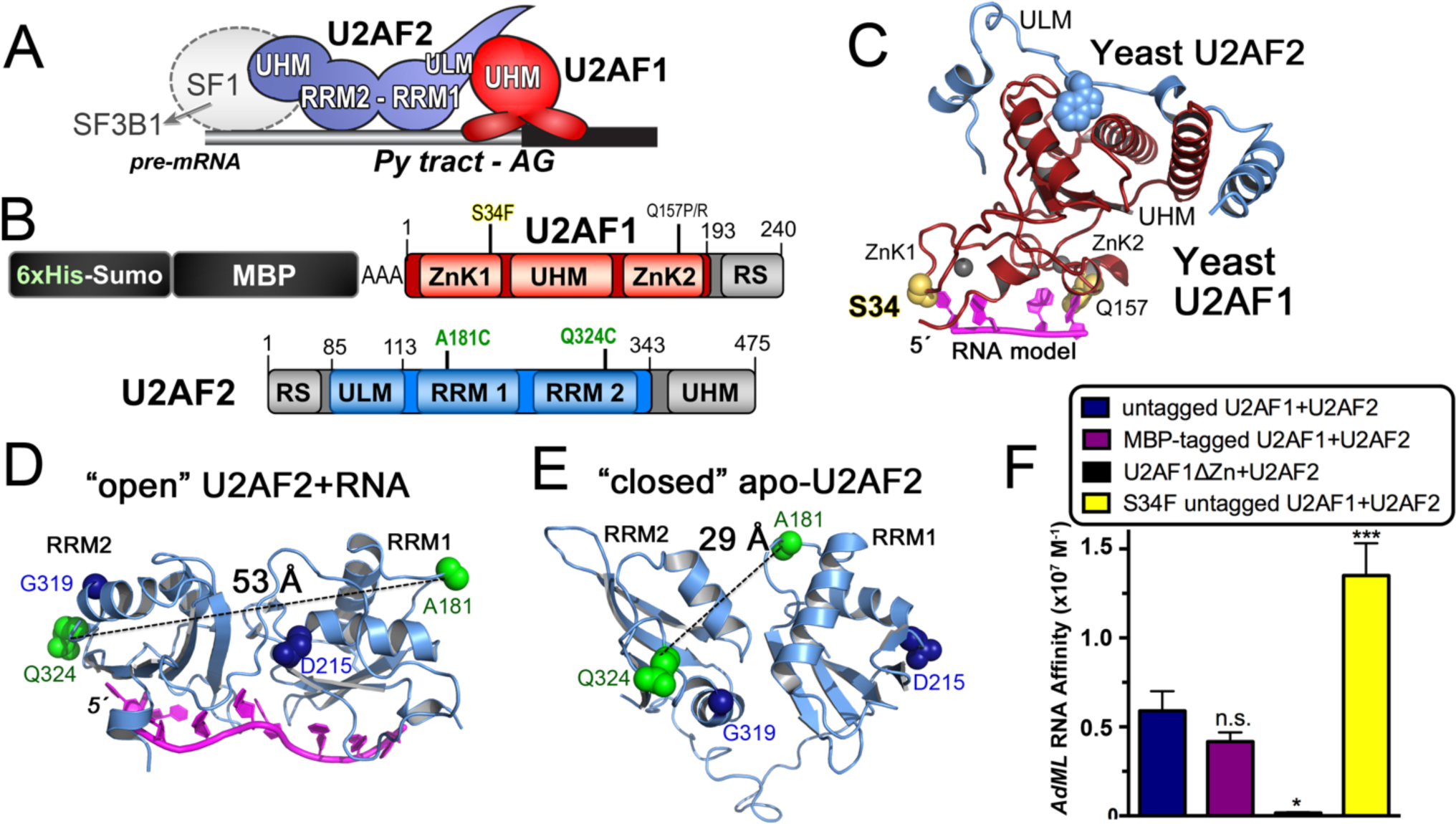
Overview of U2AF1 and U2AF2 domains, structures and labeling strategy. **(A)** U2AF1 and U2AF2 subunits recognize the polypyrimidine (Py) and AG-dinucleotide consensus signals at 3ʹ splice sites of pre-mRNA transcripts. The SF1 subunit is thought to exchange with SF3B1 during spliceosome assembly. **(B)** U2AF1 and U2AF2 domains and constructs used for smFRET (emphasized by color). The U2AF2 domains include two RNA recognition motifs (RRM1 and RRM2) and a U2AF ligand motif (ULM). U2AF1 domains include zinc knuckles (ZnK1 and ZnK2) and a central U2AF homology motif (UHM) that mediates heterodimerization. Recurrent S34F and Q157P/R cancer-associated U2AF1 mutations are marked. For surface tethering, U2AF1 is expressed as an N-terminal fusion with 6xhistidine, sumo, triple-alanine linker, and maltose-binding protein (MBP) tags. **(C)** Structure of fission yeast U2AF1 bound to U2AF2 ULM (PDB ID 4YH8) modeled with RNA. The MDS-affected residues are yellow spheres. **(D)** The open U2AF2 RRM1-RRM2 conformation bound to poly-uridine oligonucleotide (magenta) (PDB ID 5EV4). **(E)** The closed conformation of apo-U2AF2 (PDB ID 2YH0). For **(D-E)**, the U2AF2 RRMs are labeled at unique cysteine mutations of A181 and Q324 (green spheres); the D215/G319 residues (navy spheres) promote the “closed” conformation following mutation to R. **(F)** Binding affinities of U2AF heterodimer variants for the *AdML* splice site (5ʹ-fluorescein-CUGUCCCUUUUUUUUCA<u>C</u>**AG**|CUCGCGGUUGAG, where | is the spliced junction). The fitted binding curves are given in Supplementary Figure S2. not significant (n.s.) *p*>0.05; *, *p***<**0.01; ***, *p***<**0.0005 from two-tailed unpaired t-tests with Welch’s correction of the average values from three experiments for the wild-type proteins and six experiments for the S34F mutant (comparison to untagged heterodimer) calculated in GraphPad Prism.

Acquired mutations of pre-mRNA splicing factors are a prevalent source of splice site dysregulation among hematologic malignancies and myelodysplastic syndromes (MDS) (7). U2AF1 is one of the recurrently mutated splicing factors in MDS, alongside SF3B1, SRSF2, and ZRSR2. These affected splicing factors synergize for early spliceosome assembly. Interestingly, ZRSR2 is a U2AF1-paralogue of the minor, U12-type spliceosome (8,9), whereas SRSF2 shares U2AF1/U2AF2 functions of annealing pre-mRNA consensus elements with spliceosomal RNAs, such as the U2 snRNA (10-12). Protein interactions of SF3B1 with U2AF2 contribute to U2 snRNP recruitment to the 3ʹ splice site (13,14). The cancer-associated mutations of the pre-mRNA splicing factors cluster in hotspots. Most commonly, U2AF1 acquires an S34F-encoding mutation (~80% of MDS-associated *U2AF1* mutations), or in a few cases, S34Y or Q157P/R mutations (15-18). The affected U2AF1 S34 and Q157 residues are located at putative RNA interfaces of the zinc knuckles (Figure 1B,C). The S34F mutation subtly modulates U2AF1–RNA binding and splicing in a manner that correlates with the identity of the −3 nucleotide relative to the 3ʹ splice site junction (19-21). A −3C slightly enhances binding and splicing for the S34F mutant compared to wild-type U2AF1, whereas −3U splice sites are often skipped. Less frequently, cancer-associated missense mutations of the U2AF2 subunit also occur and are located at RNA-interacting residues or RRM1/RRM2 interfaces (22).

Breakthrough studies have revealed near-atomic resolution structures of spliceosome assemblies (reviewed in (23)). However, gaps remain in currently available views of the earliest stage of splice site recognition and spliceosome recruitment. Structures are available for portions of the U2AF1 and U2AF2 subunits (Figure 1C-E). A “U2AF Homology Motif” (UHM) heterodimerization domain of U2AF1 binds an N-terminal motif of U2AF2 (24,25). Two zinc knuckles fold on the opposite surface of the U2AF1 UHM (26). The U2AF1–RNA interface has yet to be elucidated by experimental structures. Meanwhile, models and RNA binding experiments suggest that the zinc knuckle motifs comprise the major AG-recognition interface of U2AF1 (20,26,27). The U2AF2 subunit recognizes the Py tract signal via tandem RNA recognition motifs (RRM1 and RRM2). High resolution structures show the U2AF2 RRMs in an open, side-by-side arrangement when bound to strong Py tracts comprising contiguous uridines (28,29). The N-terminal U2AF2 RRM1 is more promiscuous than the uridine-stringent RRM2 (30,31). In the absence of RNA, a closed conformation masks the typical RNA binding surface of RRM1, leaving only RRM2 available (29).

The U2AF2 RRMs adopt a range of associated and detached conformations when studied in solution by single molecule Förster resonance energy transfer (smFRET) and small-angle X-ray scattering (28,30,32,33). Conformational selection contributes to Py tract recognition by the isolated U2AF2 subunit, whereby preferential RNA association shifts the U2AF2 population towards the open inter-RRM arrangement. Heterodimerization of U2AF2 with a truncated U2AF1 UHM that lacks the zinc knuckle RNA interface also promotes the open U2AF2 conformation (33). Yet, whether intact U2AF1 or its recurrent S34F mutation influence U2AF2 conformations for splice site readout remains unknown.

In a step towards addressing this question, we explored the influence of nearly full length U2AF1, its cancer-associated S34F mutation, and diverse splice sites on the U2AF2 inter-RRM dynamics using smFRET. In the absence of RNA, heterodimerization with intact U2AF1 stabilizes a high FRET value of U2AF2 that corresponds to the closed inter-RRM conformation. A strong, uridine-rich splice site switches U2AF2 to a lower FRET state consistent with the open conformation.

Remarkably, a significant population of the U2AF2 conformations remain in the closed state in complexes of the heterodimer with weak, uridine-poor splice sites. This conformational difference could contribute to a conditional requirement for U2AF1 to promote splicing of degenerate splice sites. The U2AF1 S34F-mutant alters the distribution of U2AF2 conformations for a subset of weak splice site complexes, which suggests a new structural contributor to dysregulated gene expression during MDS.

## MATERIALS AND METHODS

### Protein expression and purification

For FRET experiments with the heterodimer, the wild-type and S34F-mutant U2AF1 boundaries included residues 1-189 of Refseq NP_006749 with a C-terminal R189A mutation to enhance soluble protein expression. The U2AF1 construct was fused at the N-terminus with a 6xHis-Sumo-maltose binding protein (MBP) tag in a modified pCDF-1b vector. The U2AF2 subunit included residues 85-342 of NCBI RefSeq NP_009210 fused at the N-terminus with a PreScission protease cleavable glutathione-S-transferase (GST) tag using the pGEX6p-1 vector. The U2AF2 boundaries excluded the RS domain and a cysteine-rich C-terminal domain that binds a branch site recognition factor, SF1. The endogenous U2AF2 cysteine was replaced with a natural alanine variation (C305A), and single A181C and Q324C mutations for fluorophore attachment were introduced at locations intended to maximize FRET differences between the open and closed U2AF2 conformations as described (28). The 6xHis-Sumo-MBP-U2AF1 and GST-U2AF2 proteins were expressed separately in *Escherichia coli* and purified respectively by HisTrap and GST-affinity chromatography. Prior to mixing the subunits, Superdex-75 size exclusion chromatography (SEC) was used to pre-separate the 6xHis-Sumo-U2AF1 protein from unfolded aggregates that form during over-expression of U2AF1 in the absence of U2AF2. The GST tag was cleaved and separated from U2AF2 by a heparin-affinity chromatography. The D215R-D319R-mutant U2AF2 was prepared in a similar manner.

For FRET experiments with the isolated U2AF2 subunit, U2AF2 (residues 113-342) was prepared with 6xHis and T7 tags as described (28). The minimal U2AF1-binding site (ULM, residues 85-112) was excluded from this U2AF2 construct to avoid nonspecific binding to the surface.

For RNA binding experiments, the two human U2AF subunits were co-expressed in *E. coli* and purified sequentially by MBP-affinity and GST-affinity chromatography as described (21). The GST tag was cleaved and then removed by two rounds of subtractive GST affinity chromatography. Lastly, the MBP-fused heterodimer was separated from minor aggregates by Superdex-200 SEC. For comparison with untagged U2AF1–U2AF2, the MBP was cleaved from the U2AF1 subunit using Tev protease and separated during SEC.

The fission yeast (Sp)U2AF1 and SpU2AF2 counterparts were co-expressed in *E. coli* as respective N-terminal 6xHis-GB1(PreScission site) and GST(Tev site)-fusion proteins from modified pCDF-1b and pGEX4T-2 vectors. The heterodimer was purified by sequential HisTrap and GST-affinity chromatography. The GST tag was cleaved during dialysis with Tev protease then removed by a second step of HisTrap affinity. The 6xHisGB1 tag was removed next by overnight incubation with PreScission protease, which was removed by a GST-subtractive step due to overlap in SEC. The final SpU2AF heterodimer was purified by Superdex-200 SEC.

The RNA oligonucleotides of the indicated sequences were purchased deprotected, desalted and HPLC-purified from Dharmacon, Inc.

### Sample preparation for smFRET

To remove sulfhydryl reducing agents prior to labeling, the U2AF2 subunit was diluted into 0.01% Nikkol and then dialyzed into 500 mM NaCl, 25 mM HEPES pH 7.0, 3% glycerol, 0.2 mM TCEP. The cyanine (Cy)3-maleimide (Combinix, Inc.) and Cy5-maleimide were resuspended in DMSO and mixed in an equimolar ratio immediately prior to use, then added at a 20:1 molar ratio of each dye to U2AF2 protein. The labeling reaction was incubated in the dark at room temperature for two hours and then quenched by addition of 10 mM DTT. After overnight dialysis to remove the excess dye, labeling efficiencies of >60% for each dye were calculated using the respective Cy3 and Cy5 extinction coefficients at 550 nm and 650 nm (ε^Cy3^ 150,000 M^−1^ cm^−1^, ε^Cy5^ 250,000 M^−1^ cm^−1^). The labeled U2AF2 concentration was estimated from the calculated protein extinction coefficient (17,420 M^−1^ cm^−1^) following correction for the Cy3/Cy5 absorbances at 280 nm as we have described (28). The labeled U2AF2 was mixed with stoichiometric 6xHis-Sumo-MBP-U2AF1 and the salt concentration was reduced to 150 mM NaCl, 25 mM Hepes pH 7.0, 3% glycerol, 0.2 mM TCEP by sequential dilutions with lower salt buffer. Lastly, the labeled complex was purified by Superdex-200 SEC in this buffer (**Supplementary Figure S1**). A mixture of Cy3/Cy5 fluorophores is expected at each U2AF2 site.

### Single molecule FRET data acquisition and analysis

Measurements were carried out at room temperature in 25 mM Hepes pH 7.0, 150 mM KCl, 0.01% Nikkol and 0.2 mM TCEP, 6 mM β-mercaptoethanol, 1.5 mM Trolox (to eliminate Cy5 blinking), and an oxygen-scavenging system (0.8 mg/mL glucose oxidase, 0.625% glucose, 0.02 mg/mL catalase). The sample chamber was assembled from quartz microscope slides and glass cover slips coated with a mixture of m-PEG and biotin-PEG and pre-treated with neutravidin (0.2 mg/mL). The U2AF2^Cy3/Cy5^ subunit concentration was similar in isolation and as in the heterodimer (5 nM). Excess U2AF1 (1000 nM) was mixed with the purified U2AF1– U2AF^Cy3/Cy5^ heterodimer (5 nM) before addition to the sample chamber and excess U2AF1 also was included in the imaging buffer to ensure heterodimer formation. For surface tethering of the proteins *via* 6xHis tags on the U2AF2 subunit, the sample chamber was pre-incubated with 50 nM biotinylated (Ni^+2^)NTA resin (AnaSpec, Inc.). For surface tethering of the RNA sites, RNA splice sites were synthesized as 3ʹ-conjugates with an 18-atom PEG spacer linked to a DNA oligonucleotide d(GTGCCAGCATATTTGTCGAAG) and then pre-annealed with a complementary biotinyl-DNA oligonucleotide prior to incubation (at 10 nM) with the sample chamber.

An Olympus IX71 inverted microscope, equipped with a UPlanApo 60x/1.20w objective lens was used for smFRET measurements. Cy3 and Cy5 fluorophores were excited with 532 and 642 nm lasers (Spectra-Physics). Total internal reflection (TIR) was obtained by a quartz prism (ESKMA Optics). The fluorescence emission was split into Cy3 and Cy5 fluorescence using a dual view imaging system DV2 (Photometrics) equipped with a 630 nm dichroic mirror and detected with an Andor iXon þ EMCCD camera. Movies were recorded using the Single software (courtesy of Prof. Taekjip Ha at John Hopkins U., http://ha.med.jhmi.edu/resources/) with an exposure time of 100 ms. We typically acquired up to five 5-min movies while imaging different sections of the slide for each sample. Before each measurement, we confirmed that non-specific binding was virtually absent by imaging U2AF2^Cy3/Cy5^ added to the slide in the absence of neutravidin.

Data sets were processed using scripts from http://ha.med.jhmi.edu/resources/ (Prof. T. Ha) with IDL and MATLAB software. Apparent FRET efficiencies (E_*app*_) were calculated using the equation *E*_*app*_= *I*_*Cy5*_/(*I*_*Cy5*_+*I*_*Cy3*_); where *I*_*Cy3*_ and *I*_*Cy5*_ are the respective intensities of Cy3 and Cy5, using a MATLAB script generously provided by Prof. Peter Cornish (U. Missouri, Columbia). The FRET distribution histograms were built from traces that showed single photobleaching steps for Cy3 and Cy5 and/or anti-correlated events of donor and acceptor intensities. Anti-correlated changes in donor and acceptor intensities with constant sum of intensities indicated the presence of an energy transfer in single molecules labelled with one donor and one acceptor dye. All histograms were smoothed with a five-point window and plotted using Origin software (OriginLab Co.).

### Fluorescence anisotropy equilibrium RNA affinity measurements

The apparent equilibrium binding affinities were fit from changes in the fluorescence anisotropies of 5ʹ-fluorescein-labeled RNA oligonucleotides during titration with the indicated protein complexes as described (34). The binding buffers included 25 mM HEPES pH 6.8 and 0.2 U/μL SUPERase-In^TM^ RNAse inhibitor for all samples. The binding buffer of the human heterodimers also included 150 mM NaCl, 0.5 mM TCEP, 3% glycerol, and 20 μM ZnCl_2_. The binding buffer of the fission yeast heterodimers also included 100 mM NaCl and 0.2 mM TCEP.

### Measurement of biomolecular interaction rates

The on/off rates of the wild-type or S34F-mutated fission yeast U2AF heterodimer interacting with RNA were measured by fitting surface plasmon resonance changes detected using a BIAcore T200 instrument. A C1 sensorchip (GE Healthcare) pre-washed with glycine pH 12/0.3% Triton X-100 was amine-coupled with neutravidin at pH 5.5 and then conditioned with three injections of 1 M NaCl/50 mM NaOH. The sensorchip was equilibrated with degassed running buffer of 100 mM NaCl, 10 mM HEPES pH 6.8, 0.005% Tween 20, 0.2 mM TCEP, prepared with nuclease-free water (Sigma-Aldrich). The 5ʹ-biotinyl-RNA oligonucleotide (5ʹ-Bi.UAAGAAAUACUAAAUUAAUUUC(U/C)AG | AAAAGAGUCU for *DEK −3U/C*) was coupled to the sensorchip to a level of approximately 40 RU. Interaction rate data were collected at a flow rate of 50 μL min^−1^. Before data collection, 1 μM protein was injected twice without regeneration to reduce nonspecific binding in subsequent experiments. The kinetic experiments included 90 s sample contact time, 180 s dissociation, 30 s stabilization in running buffer. Between sample injections at the concentrations indicated in Figure 8, the surface was regenerated by treatment with 10 mM glycine pH 2.5 for 30 s followed by 60 s stabilization in running buffer. Since RNA leached during regeneration, the surface was re-treated with 5ʹ-biotinyl-RNA following regeneration and prior to the next injection.

**Figure 8.**
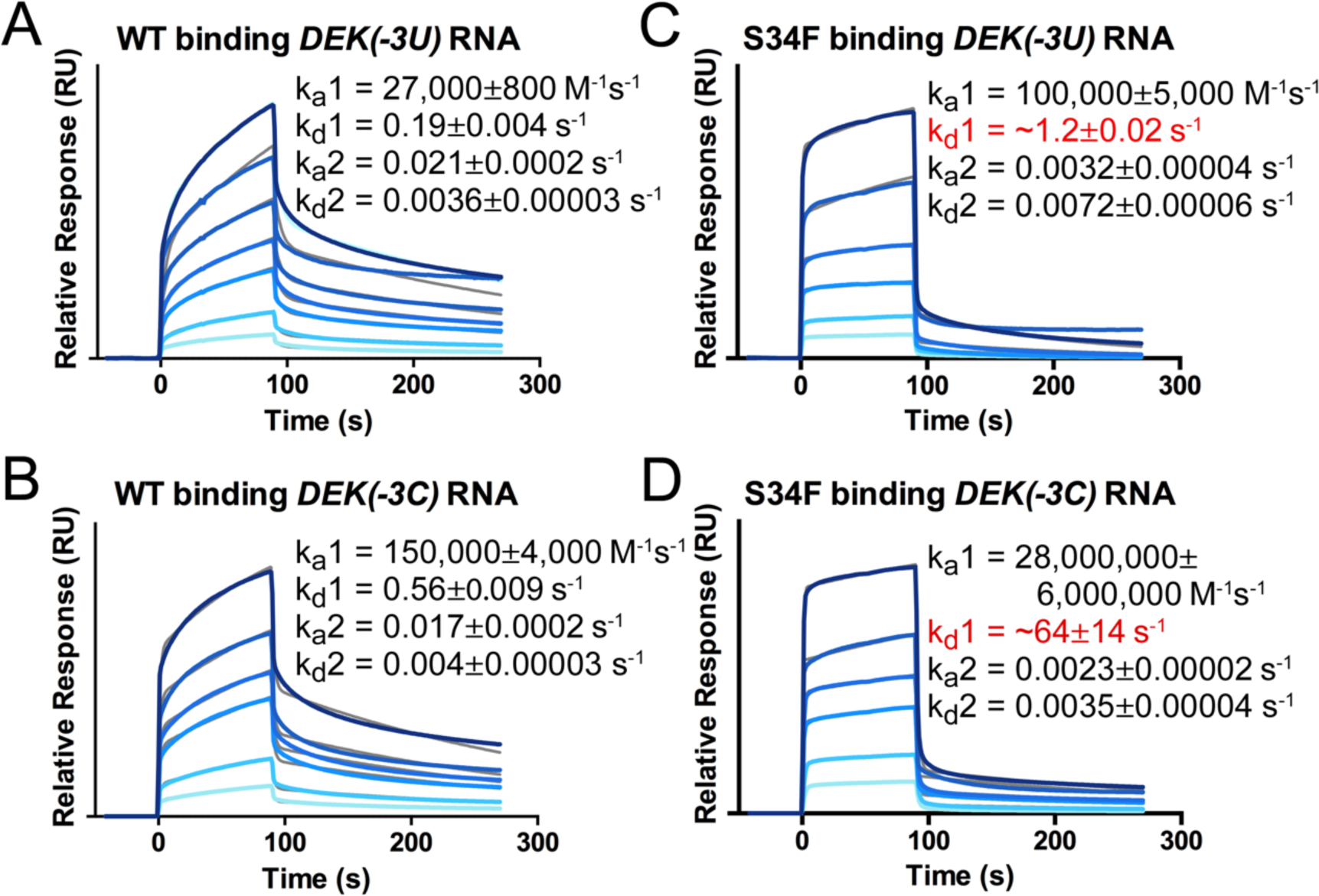
The kinetics of *Schizosaccharomyces pombe* (Sp)U2AF1–U2AF2 heterodimer binding *DEK(−3U)* or *DEK(−3C)* splice site RNAs are two-state and altered by the S34F substitution. **(A)** wild-type (WT) SpU2AF1–U2AF2 binding *DEK(−3U)* RNA; **(B)** S34F SpU2AF1–U2AF2 binding *DEK(−3U)* RNA; **(C)** WT SpU2AF1–U2AF2 binding *DEK(−3C)* RNA; **(D)** S34F SpU2AF1–U2AF2 binding *DEK(−3C)* RNA. Global two-state fits (gray) are overlaid on the experimental sensorgrams, which are colored in a gradient from light to navy blue over the range of protein concentrations (125, 250, 500, 750, 1000, 2000 nM). The biotinyl-RNA sites are immobilized on neutravidin sensorchips. Red font indicates values that exceed the reliable detection limits of the BIAcore T200 (1E+03 < k_a_1 < 3E+09 M^−1^ s^−1^ and 1E-05 < k_d_1 < 1E00 s^−1^).

## RESULTS

### U2AF1 stabilizes a high FRET conformation of U2AF2^Cy3/Cy5^

We leveraged our prior fluorophore sites in the U2AF2 RRMs (28) and methods to prepare nearly full length human U2AF1 (19,21) to investigate the influence of U2AF1 on the U2AF2 inter-RRM dynamics using smFRET. We labeled unique cysteines introduced in the N-terminal RRM1 (A181C) and C-terminal RRM2 (Q324C) of U2AF2 (Figure 1B) with an equimolar mixture of Cy3 and Cy5 fluorophores as described (28). These fluorophore positions were chosen to maximize the expected differences in FRET efficiencies between the open (lower FRET) and closed (higher FRET) U2AF2 conformations (Figure 1D-E). To study the U2AF heterodimer, we lengthened the U2AF2 construct to include the U2AF1 binding site (U2AF ligand motif, ULM), mixed labeled U2AF2^Cy3/Cy5^ with the unlabeled U2AF1 subunit, and purified the heterodimer by size exclusion chromatography (**Supplementary Figure S1**). An N-terminal 6xHis-Sumo-MBP tag tethered U2AF1 with bound U2AF2^Cy3/Cy5^ to a biotin-NTA/Ni^+2^ slide (Figure 1B, Figure 2A). The MBP tag reduced nonspecific interactions with the slide surface and had no detectable effect on the RNA binding affinity of the heterodimer (Figure 1F, **Supplementary Figure S2**). Moreover, we corroborated that the U2AF1 zinc knuckles are actively folded by showing that these motifs contribute 25-fold to the heterodimer–RNA binding affinity. By tethering the labeled U2AF2^Cy3/Cy5^ subunit *via* U2AF1 and including excess unlabeled U2AF1 in the imaging buffer (Methods), we ensured imaging of only heterodimeric U2AF1-–U2AF2^Cy3/Cy5^ complexes. For a well-controlled comparison, we also replicated smFRET experiments with the U2AF2^Cy3/Cy5^ subunit alone, which we tethered *via* an N-terminal 6xHis-tag as described (28) (Figure 2D).

**Figure 2.**
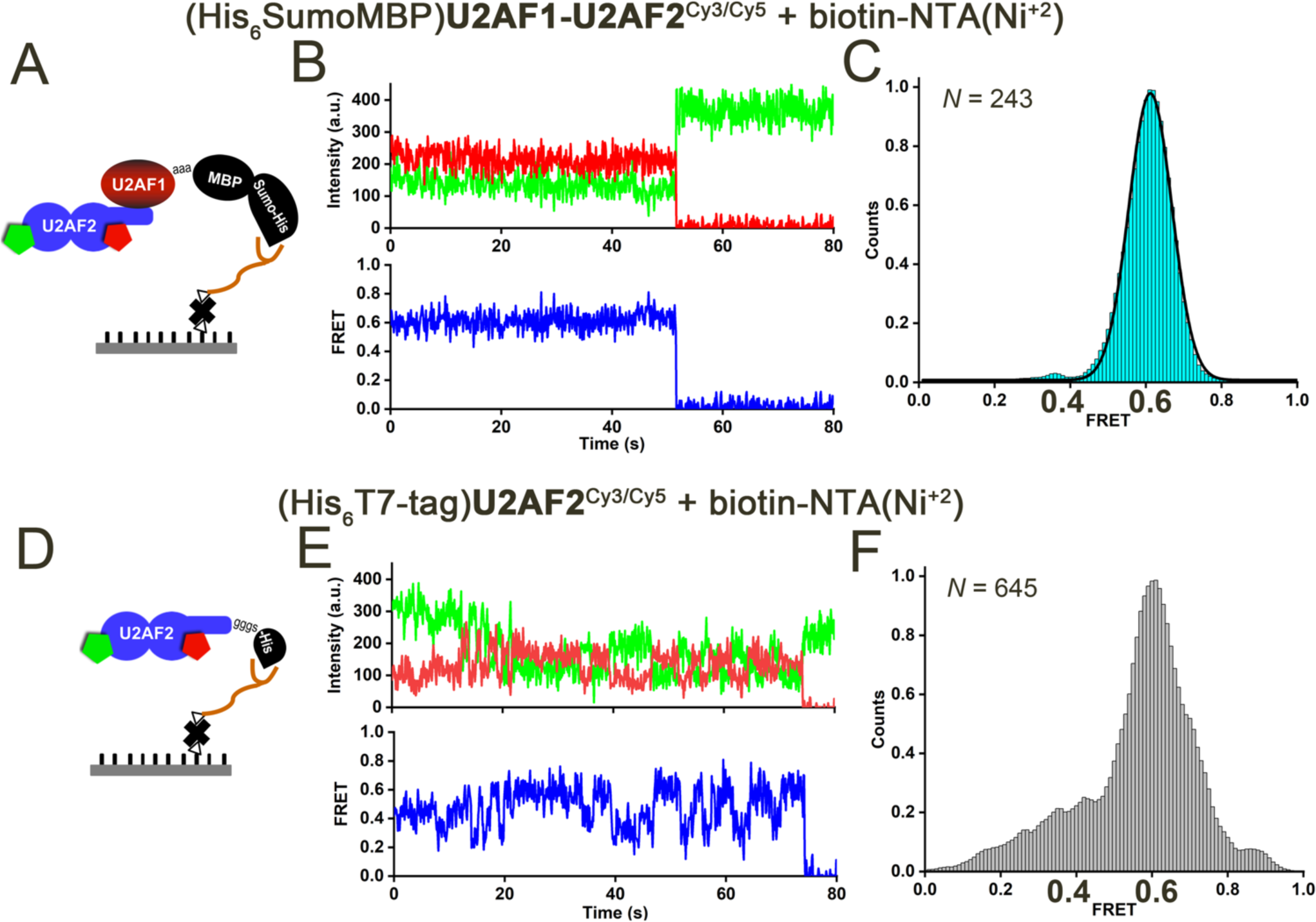
U2AF1 stabilizes a U2AF2 conformation with a high FRET value in the absence of RNA. **(A)** Scheme for U2AF1–U2AF2^Cy3/Cy5^ immobilization. The histidine-tagged U2AF1 (purified as a heterodimer with U2AF2^Cy3/Cy5^) was immobilized *via* biotinyl-NTA(Ni^+2^) on a neutravidin-coated surface. A mixture of Cy3/Cy5 is present at each labeled site. **(D)** The histidine-tagged U2AF2 subunit is immobilized directly as described (28). **(B)** Representative smFRET trace of the U2AF1– U2AF2^Cy3/Cy5^ heterodimer in absence of RNA show fluorescence intensities from Cy3 (green) and Cy5 (red) and the calculated apparent FRET efficiency (blue). The majority of U2AF1-bound U2AF2^Cy3/Cy5^ traces lack fluctuations and show predominate FRET values of ~0.6. **(E)** Representative traces of the U2AF2 subunit alone, which is dynamic compared to the U2AF1-bound heterodimer. **(C, F)** Histograms compiled from *N* FRET traces showing the distribution of FRET values for (**C**) U2AF1– U2AF2^Cy3/Cy5^ heterodimer (cyan) or (**F**) U2AF2^Cy3/Cy5^-only (gray). A black line represents the Gaussian fit of the U2AF1–U2AF2^Cy3/Cy5^ histogram in (E).

The presence of the U2AF1 subunit stabilized a high FRET state of U2AF2^Cy3/Cy5^. As previously observed using smFRET or small-angle X-ray scattering (28,32), the apo-U2AF2^Cy3/Cy5^ FRET distribution histogram was broad (Figure 2F). Approximately 30% of the apo-U2AF2^Cy3/Cy5^ smFRET traces showed transitions between different FRET values (Figure 2E). By contrast, nearly all of the U2AF1–U2AF2^Cy3/Cy5^ traces (95%) showed stable FRET values of approximately 0.6 (Figure 2B). Accordingly, the FRET distribution histogram built from hundreds of U2AF1–U2AF2^Cy3/Cy5^ traces was narrowly centered at a FRET value of 0.6. We concluded that heterodimerization with the U2AF1 subunit stabilized a particular U2AF2 inter-RRM1/RRM2 arrangement with a characteristic high FRET value.

### A uridine-rich Py tract switches U2AF2^Cy3/Cy5^ to a U2AF1-stabilized lower FRET conformation

We then used smFRET to investigate the influence on U2AF2 conformations of binding to a strong, uridine-rich splice site from the adenovirus major late promoter prototype (*AdML*) (sequence given in Figure 3A). Consistent with prior observations (28), addition of the isolated U2AF2^Cy3/Cy5^ subunit to the slide-tethered *AdML* splice site selectively increases a fraction of molecules with 0.4 FRET values out of an otherwise broad FRET distribution (**Supplementary Figure S3A-B**). Remarkably for the tethered U2AF1–U2AF2^Cy3/Cy5^ heterodimer, addition of the *AdML* RNA switched the predominant 0.6 FRET state of the FRET distribution histogram to a narrow peak centered at an approximately 0.4 FRET value (Figure 3B-C, **Supplementary Figure S4**). A similar 0.4 FRET state resulted from analysis of a reverse immobilization strategy, in which untethered U2AF1–U2AF2^Cy3/Cy5^ was added to AdML RNA attached to the slide (Figure 3D-E). The AdML RNA-bound heterodimer complex was extremely stable; indeed, 99% of the traces showed 0.4 FRET until photobleaching. We concluded that a large *AdML* RNA-dependent change in the U2AF1-bound U2AF2^Cy3/Cy5^ conformation occurred and was independent of possible structural heterogeneity introduced by RNA-free proteins or tethering to the surface.

**Figure 3.**
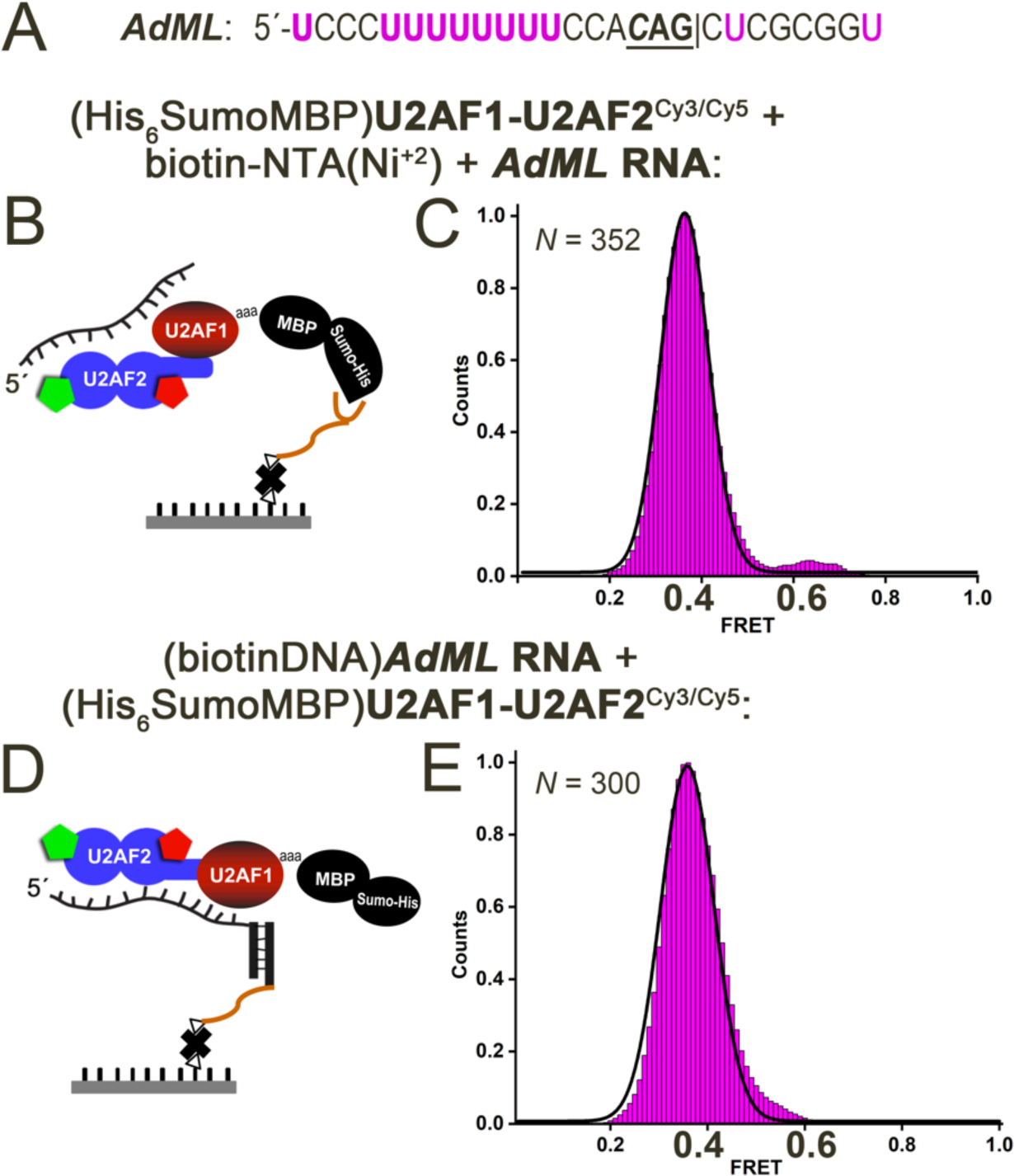
A uridine-rich, strong splice site RNA (*AdML*) switches the U2AF1–U2AF2^Cy3/Cy5^ heterodimer to a U2AF2 conformation corresponding to a lower FRET state. **(A)** Sequence of the *AdML* splice site oligonucleotide: “|” spliced junction; three preceding nucleotides are bold and the −3 nucleotide is italicized. **(B)** Scheme for U2AF1–U2AF2^Cy3/Cy5^ protein immobilization and RNA addition, which is analogous to Figure 2A. **(C)** Histogram showing the distribution of FRET values for tethered U2AF1–U2AF2^Cy3/Cy5^ following addition of *AdML* RNA. **(D)** Scheme for RNA immobilization and U2AF1–U2AF2^Cy3/Cy5^ protein addition. An RNA splice site-(PEG)-DNA conjugate was annealed with a biotin-DNA oligonucleotide and immobilized on a neutravidin-coated biotin-Peg surface. **(E)** Histogram showing the distribution of FRET values for tethered *AdML* RNA bound to U2AF1–U2AF2^Cy3/Cy5^. Black lines represent the Gaussian fits of the histograms. *N*, total number of compiled single-molecule traces.

### A D215R/G319R mutant indicates that the high FRET state corresponds to closed U2AF2

To investigate whether the 0.6 FRET state of U2AF2^Cy3/Cy5^ corresponds to the well-characterized closed conformation, we introduced a D215R/G319R double-mutation that strengthens the closed U2AF2 conformation by electrostatic complementarity (29) (Figure 1D-E). As expected, the already 0.6 FRET conformation of U2AF1–U2AF2^Cy3/Cy5^ appeared unchanged in the RNA-free histogram (Figure 4A). We then tested adding U2AF1–U2AF2^Cy3/Cy5^ to tethered *AdML* oligonucleotide, which ensured that only RNA-bound complexes were analyzed. Rather than shifting the entire U2AF2^Cy3/Cy5^ population to the lower 0.4 FRET value as observed for the wild-type heterodimer, approximately half of the conformational ensemble of the D215R/G319R mutant remained in the 0.6 FRET state (Figure 4B). This result supported our assignment of the 0.6 FRET state as the closed U2AF2 conformation, whereas the 0.4 FRET state is consistent with the open arrangement of U2AF RRM1/RRM2.

**Figure 4.**
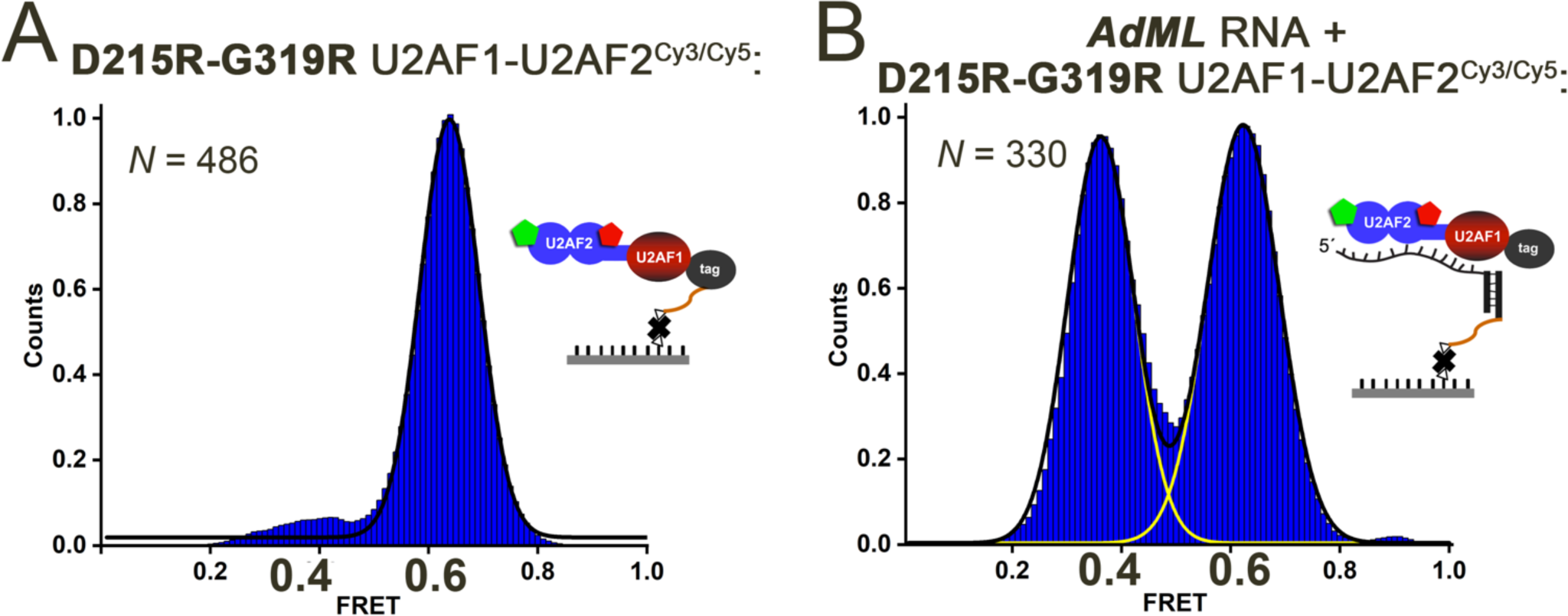
A D215R/G319R variant that is known to stabilize the closed U2AF2 conformation (29) promotes U2AF2^Cy3/Cy5^ conformations corresponding to the ~0.6 FRET state. Histograms of **(A)** tethered U2AF1–D215R/G319R U2AF2^Cy3/Cy5^ in the absence of RNA or **(B)** tethered *AdML* RNA in the presence of U2AF1–D215R/G319R U2AF2^Cy3/Cy5^. The immobilization strategies are inset. Black and yellow lines indicate the respective summed or individual Gaussian fits.

### Uridine-poor Py tracts bind open and closed U2AF1-stabilized U2AF2^Cy3/Cy5^ conformations

Considering that the U2AF1 subunit is required for splicing a subset of weak splice sites with short, degenerate Py tracts, we next used smFRET to probe the influence of weak splice sites on the U2AF2 conformation in the U2AF1–U2AF2^Cy3/Cy5^ heterodimer (Figure 5). We started with a prototype of U2AF1-dependent splice sites from the immunoglobulin M gene (*IgM*) (35), and compared splice sites that have been shown to be affected by the MDS-associated S34F mutation (*DEK*, *CASP8*, and *FMR1*) (19,21). The binding affinities of the heterodimer for these degenerate splice sites were very weak (e.g. K_D_ >3 μM for *DEK* splice site in **Supplementary Figure S5**, which is 100-fold weaker than the ternary U2AF1–U2AF2–SF1 complex (19)). As such, we focused on the immobilization strategy of adding U2AF1–U2AF2^Cy3/Cy5^ heterodimer to the tethered RNAs (Figure 3D), which avoids introducing heterogeneity from RNA-free proteins. We also compared analogous smFRET experiments for the isolated U2AF2^Cy3/Cy5^ subunit bound to representative *IgM* and *DEK* RNAs and found broader but qualitatively similar histograms (**Supplementary Figure S3C-E**). Unlike the complete shift to a lower 0.4 FRET state observed for the heterodimer bound to strong *AdML* site, the *IgM* prototype of U2AF1-dependent splice sites binds most of the U2AF2^Cy3/Cy5^ molecules in the closed, 0.6 FRET conformation (~70% of traces) (Figure 5A). The remaining population of traces (~30%) shifts U2AF2^Cy3/Cy5^ to the 0.4 FRET state that are likely to correspond to the open conformation. Likewise, the smFRET distribution histograms built from hundreds of U2AF1– U2AF2^Cy3/Cy5^ traces bound to uridine-poor *DEK*, *CASP8*, or *FMR1* splice sites showed populations of both 0.4 and 0.6 FRET traces at various ratios (Figure 5B-D). We concluded that populations of both open and closed U2AF2 conformations bind to splice sites with degenerate Py tracts.

**Figure 5.**
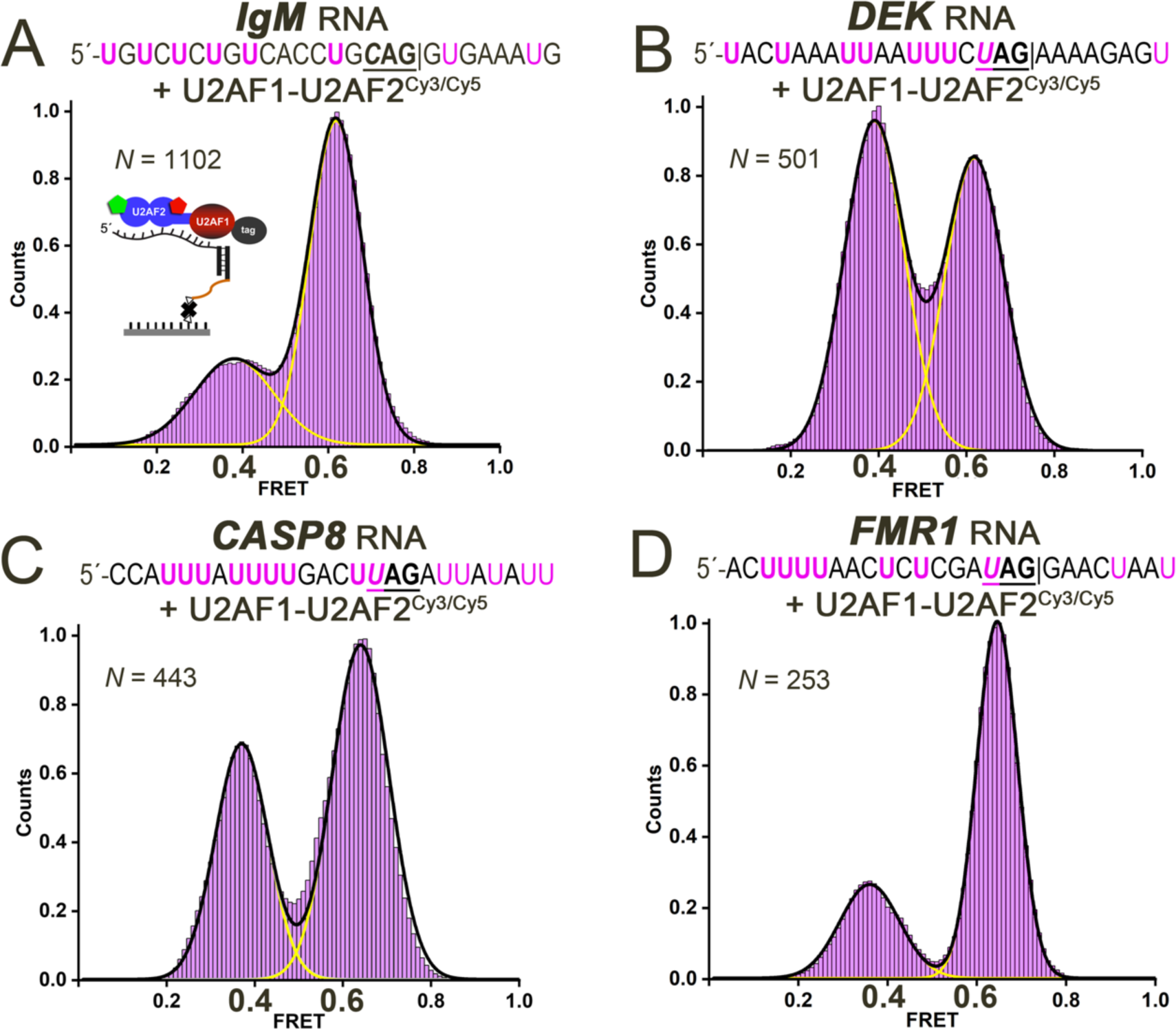
Binding to weak, uridine-poor splice sites alters the equilibrium between closed (0.6 FRET) and open (0.4 FRET) conformations of U2AF1–U2AF2^Cy3/Cy5^. Histograms showing the distribution of FRET values for the respective **(A)** *IgM*, **(B)** *DEK* (also called *DEK(−3U)* in Figure 7), **(C)** *CASP8* or **(D)** *FMR1-*tethered complexes of U2AF1–U2AF2^Cy3/Cy5^ (lavender). The RNA is immobilized ensure that RNA-bound states are detected (scheme in panel A). The RNA sequences are given above each panel. Black and yellow lines indicate the respective summed or individual Gaussian fits.

Whereas apparent transitions between different FRET values were extremely rare for the U2AF1–U2AF2^Cy3/Cy5^ heterodimer bound to the uridine-rich *AdML* splice site, the U2AF1– U2AF2^Cy3/Cy5^ complexes with uridine-poor splice sites showed multiple spontaneous transitions between 0.4 and 0.6 FRET for 10 - 20% of traces. These characteristics of the heterodimer differed from the typically single, directional changes from ~0.6 to ~0.4 FRET that we had previously observed for the isolated U2AF2^Cy3/Cy5^ subunit binding to the *AdML* splice site (28). The single irreversible transitions from 0.6 to 0.4 FRET likely indicates that the uridine-rich *AdML* splice site stabilizes the open U2AF2 conformation. In contrast, the fluctuations of U2AF2^Cy3/Cy5^ between 0.4 and 0.6 FRET reinforces that both open and closed conformations are compatible with binding to uridine-poor sites.

### The S34F mutation of U2AF1 modulates the U2AF2^Cy3/Cy5^ FRET distribution

The MDS-associated S34F mutation occurs at a putative U2AF1–RNA interface and could influence the conformation of the U2AF2 subunit for Py tract recognition. We used smFRET to investigate the effects of the U2AF1 S34F mutation on the U2AF2^Cy3/Cy5^ conformations, first in the absence of RNA and then bound to the uridine-rich *AdML* or various uridine-poor splice sites (Figure 6). The S34F mutation subtly promoted the closed, 0.6 FRET state of U2AF2^Cy3/Cy5^ in the context of the RNA-free S34F mutant heterodimer (Figure 6A). Similarly, subtle or no differences were detected in the FRET distributions of the S34F mutant compared to wild-type U2AF1–U2AF2^Cy3/Cy5^ complexes with *AdML*, *IgM*, *DEK*, or *CASP8* splice site RNAs (Figure 6B-E). The differences between the wild-type and S34F mutant U2AF1–U2AF2^Cy3/Cy5^ conformations were most evident for the complex with the *FMR1* splice site. Most of the *FMR1*-bound wild-type heterodimers were detected in the closed, 0.6 FRET conformation of U2AF2^Cy3/Cy5^ (74% of traces), whereas introduction of the S34F mutant switches U2AF2^Cy3/Cy5^ to the open, 0.4 FRET conformation (66% of traces) (Figure 6F). The reason for this reproducible, RNA sequence-dependent difference is not certain. One conjecture is that the bulky phenylalanine might slightly strain the closed U2AF2 conformation when complexed with certain splice sites, such that the population shifts towards the open state when upstream U-rich RNA sequences are available for binding (as for *FMR1*).

**Figure 6.**
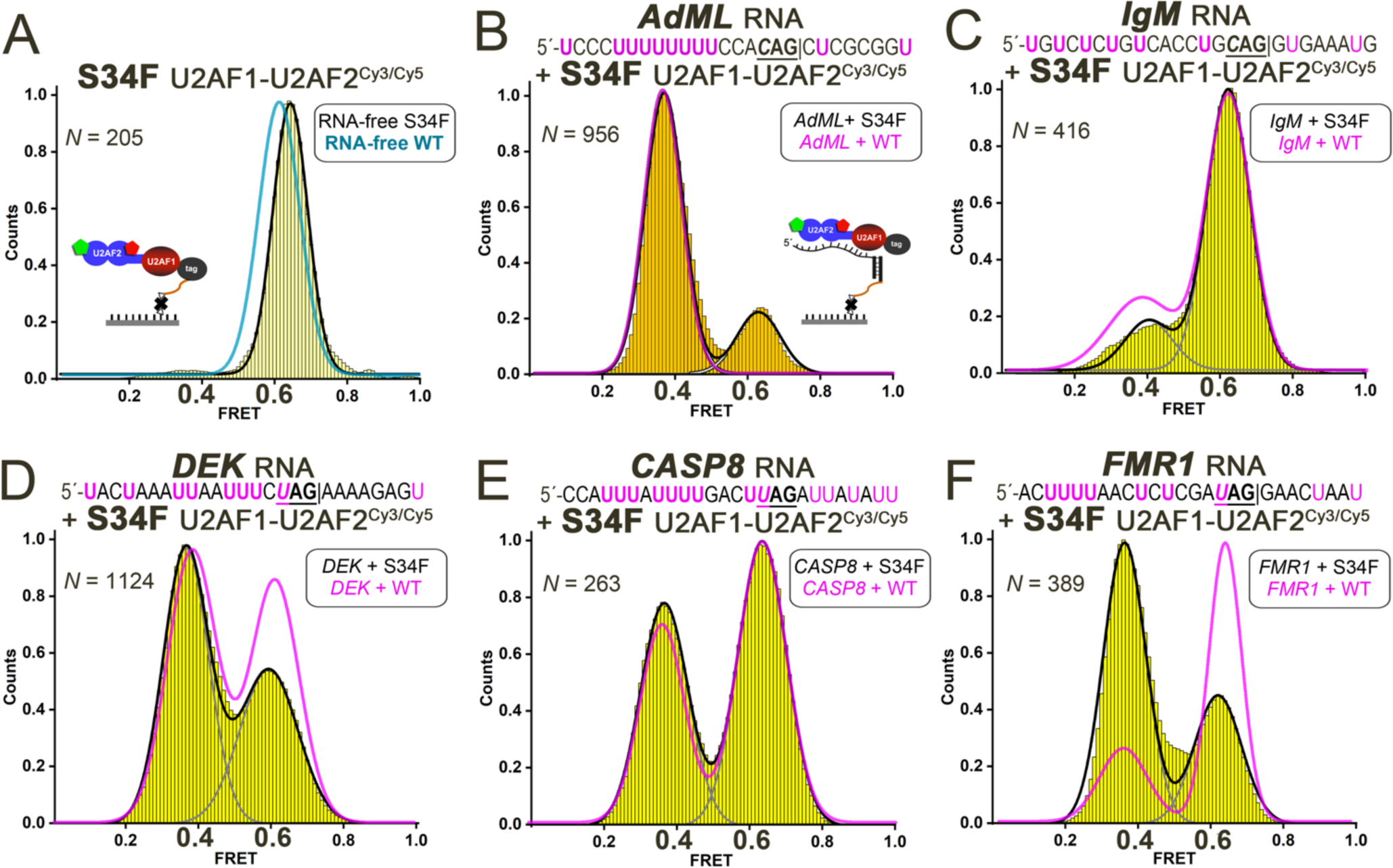
Influence of the MDS-associated S34F U2AF1 (yellow) on the U2AF1–U2AF2^Cy3/Cy5^ FRET distributions. Histograms of S34F-mutated U2AF1–U2AF2^Cy3/Cy5^ heterodimer either **(A)** in the absence of RNA, or **(B-F)** bound to tethered RNAs including **(B)** *AdML*, **(C)** *IgM*, **(D)** *DEK* (also called *DEK(−3U)* in Figure 7), **(E)** *CASP8*, **(F)** *FMR1.* The RNA-free protein in panel A is tethered by the His_6_-tag on U2AF1 as in Figure 2A (scheme inset). The RNA-bound proteins in panels B-F are tethered by annealing a DNA-linked splice site RNA with a biotin-DNA oligo as in Figure 3D (scheme inset in panel B). The splice site RNA sequences are given above the histograms. Black and gray lines indicate the respective summed or individual Gaussian fits of the S34F mutant histograms (widths <0.14). The corresponding summed fits of wild-type U2AF1–U2AF2^Cy3/Cy5^ (pale cyan for RNA-free or magenta for RNA-bound) from Figures 2C, 3E and 6 are overlaid for comparison.

### U2AF1-stabilized U2AF2^Cy3/Cy5^ conformations sense the −3C/U preceding the spliced junction

Both wild-type and S34F slightly prefer splicing and binding to sites with a cytidine compared to a uridine at the −3 nucleotide preceding the spliced junction, and this preference is enhanced by the S34F mutation (19-21). Since the arrangement of the splice site signals relative to the U2AF1 and U2AF2 subunits are unknown, it is possible that U2AF2 contacts the −3C/U nucleotide and binding differences are indirectly influenced by the U2AF1 S34F mutation. We first investigated the influence of a −3C variant for the *DEK* splice site, which normally is preceded by a −3U, on the conformational ensemble of the isolated U2AF2^Cy3/Cy5^ subunit. The *DEK(−3C)* variant preferentially associated with the open U2AF2^Cy3/Cy5^ conformation, which differed from the open and closed U2AF2^Cy3/Cy5^ populations bound to the original *DEK(−3U)* RNA (**Supplementary Figure S3D-E**). As for other splice sites, the U2AF1 subunit sharpened the FRET distributions of U2AF2^Cy3/Cy5^ in the heterodimer bound to the *DEK(−3C)* RNA (Figure 7). Binding to the *DEK(−3C)* RNA increased the population of open U2AF1– U2AF2^Cy3/Cy5^ conformations compared to the original *DEK(−3U)* splice site (Figure 7A). We next extended our experiments to a −3U-substituted *IgM* splice site, which naturally is preceded by a −3C. The artificial *IgM(−3U)* nucleotide again increased the population of open U2AF1–U2AF2^Cy3/Cy5^ conformations compared to the natural *IgM(−3C)* parent sequence (Figure 7B), despite a C-to-U change for the artificial *IgM* rather than the opposite U-to-C variation of the *DEK* splice site.

**Figure 7.**
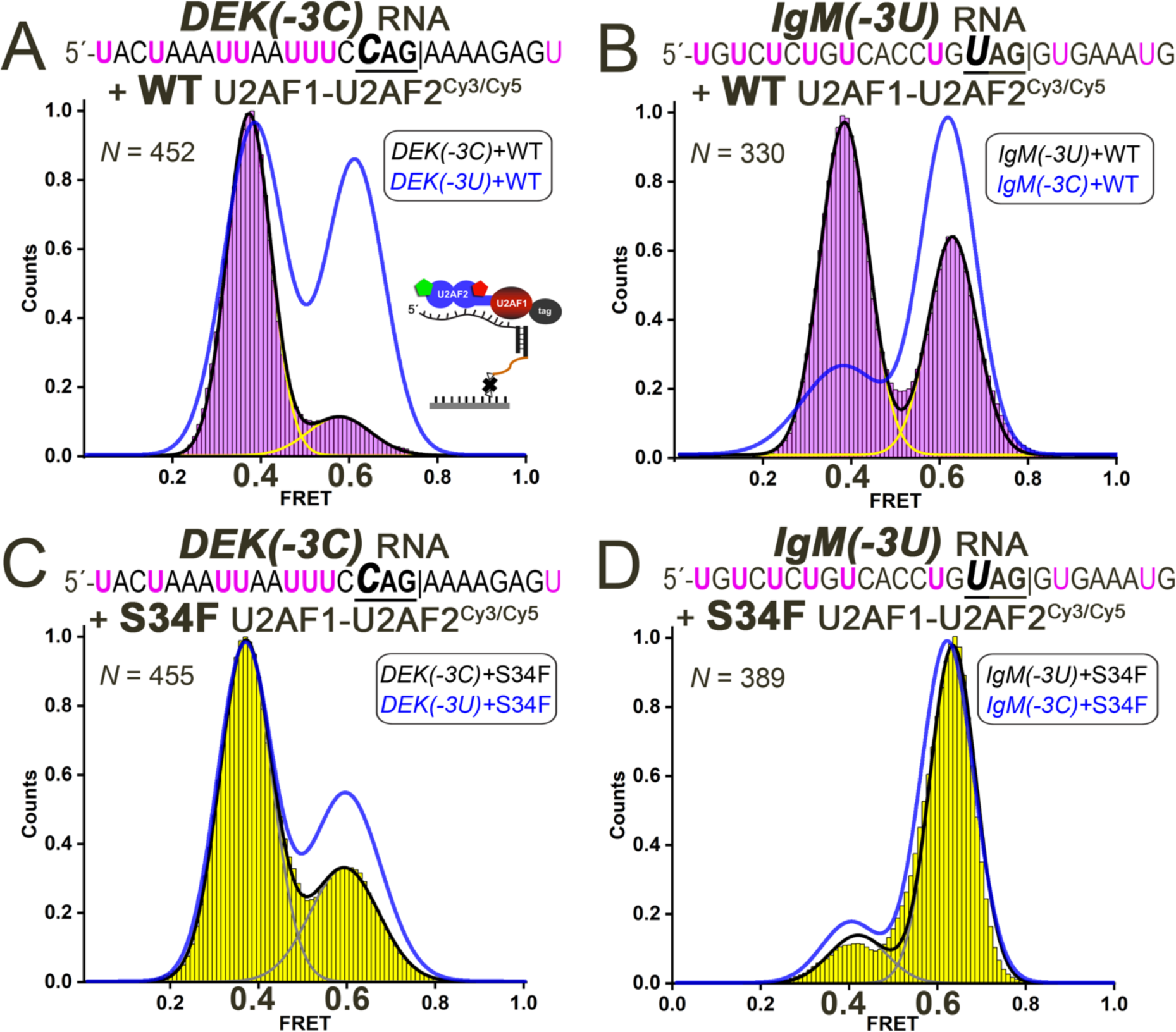
Influence of a −3C *versus* −3U nucleotide preceding the splice site junction on the FRET distributions of the wild-type (WT) and S34F mutant U2AF1–U2AF2^Cy3/Cy5^ heterodimers, including **(A)** WT U2AF1–U2AF2^Cy3/Cy5^ bound to *DEK(−3C)* RNA, **(B)** WT U2AF1–U2AF2^Cy3/Cy5^ bound to *IgM(−3U)* RNA, **(C)** S34F U2AF1–U2AF2^Cy3/Cy5^ bound to *DEK(−3C)* RNA, **(D)** S34F U2AF1– U2AF2^Cy3/Cy5^ bound to *IgM(−3U)* RNA. The ribonucleoproteins are tethered *via* the bound RNAs as inset in panel A and shown in Figure 3D. Black lines indicate the summed Gaussian fits of the histograms, and yellow or gray lines indicate the individual fits. The summed fits of the FRET histograms for the natural splice sites (*DEK(−3U)* or *IgM(−3C)*) bound to the corresponding WT or S34F mutant heterodimers are overlaid in blue for comparison.

We then compared the influence of the S34F mutation on the FRET distributions of the U2AF1–U2AF2^Cy3/Cy5^ heterodimer bound to the −3U/C variant splice sites. The differences between the *DEK(−3C)* compared to the *DEK(−3U)* splice site complexes were more subtle for the S34F mutant compared to the wild-type heterodimer (Figure 7C), although the open, ~0.4 FRET population still increased slightly for the S34F-mutant U2AF1–U2AF2^Cy3/Cy5^ bound to artificial *DEK(−3C)* RNA compared to the original splice site. For the S34F mutant heterodimer bound to *IgM(−3U)* RNA, the FRET distribution histograms remained indistinguishable compared to the *IgM(−3C)* splice site (Figure 7D). We concluded that in general, the open/closed U2AF2 conformations in the wild-type U2AF1 heterodimer normally sense the cytidine/uridine identity at the −3 position of the splice sites and are influenced by the RNA sequence of the surrounding splice site. Remarkably, the S34F mutation reduced the ability of U2AF2 to conformationally respond to the −3U/C identities of the splice sites tested here.

### Rates of a U2AF heterodimer interacting with *DEK* RNA reflect two conformational states and are rapid in the presence of the S34F mutation

The presence of two distinct U2AF conformations predicted that the kinetics for splice site RNA binding would be complex. Moreover, we hypothesized that larger differences in the interaction rates underpinned the subtle influence of the S34F mutation on the equilibrium U2AF–RNA binding affinities. To investigate these possibilities, we used Biomolecular Interaction Analysis (BIAcore) of surface plasmon resonance changes to measure the rates of a U2AF1–U2AF2 heterodimer binding to representative RNAs. Since the human homologues nonspecifically bound the carboxymethylated dextran sensorchip, we substituted a *Schizosaccharomyces pombe* (Sp) yeast heterodimer of full length U2AF1 and a U2AF2 region corresponding to the human boundaries shown in Figure 1B (77%/47% conservation between *S. pombe* and human U2AF1/U2AF2 constructs). We focused on the *DEK* splice site and its *-3C* variant, since *DEK* is a well-characterized S34F-skipped splice site (19) bound by populations of U2AF1–U2AF2^Cy3/Cy5^ in both 0.4 and 0.6 FRET states. We first checked the equilibrium RNA binding affinities of the wild-type and S34F-mutated SpU2AF heterodimer by monitoring fluorescence anisotropy changes during titration of the proteins into fluorescein-tagged *DEK* variants (**Supplementary Figure S6**). The results agreed with prior measurements for the ternary human U2AF1–U2AF2–SF1 complex (19): the fission yeast heterodimer slightly preferred binding to the *DEK(−3C)* over *DEK(−3U)* splice sites and this effect was enhanced by the S34F mutation.

We tethered biotinylated *DEK* RNA sites to neutravidin sensorchips and used a BIAcore T200 instrument to measure the kinetic rate constants of binding to the wild-type or S34F mutant SpU2AF heterodimer (Figure 8). As expected considering that two major conformations of the human homologue bound the *DEK* site, the fits of all sensorgrams were improved by two state models (up to 40-fold decrease in χ-values). Whereas the RNA interaction rates of the wild-type SpU2AF were within the reliable range of the kinetic fit, the S34F mutant interactions were extremely rapid. Indeed, the dissociation rate constants of the S34F SpU2AF2 were beyond the reliable limits of the instrument for either *DEK* variant. Altogether, the *DEK* RNA interaction kinetics of the SpU2AF heterodimer are consistent with the two major U2AF2 conformations and further reveal a large influence of the S34F mutation on the rates of formation and dissociation of the ribonucleoprotein complex.

## DISCUSSION

Our smFRET data reveals RNA-responsive switching between the open and closed conformations of U2AF2 when in the context of U2AF1–U2AF2 heterodimer (Figure 9). Although extended conformations, in which the two U2AF2 RRMs fully dissociate, may exist beyond the detectable limit of our Cy3/Cy5 FRET experiment, the narrow FRET distributions of the U2AF1–U2AF2^Cy3/Cy5^ heterodimer support the presence of only two close-packed arrangements of RRM1/RRM2. The U2AF2 conformations of the human U2AF heterodimer and its ribonucleoprotein complexes are remarkably stable and rarely show dynamic transitions in the smFRET traces. Heterodimerization with the U2AF1 subunit is necessary to structurally reinforce these distinct structural states of U2AF2. Otherwise, a continuum of less populated inter-RRM1/RRM2 distances are evident in the FRET histograms of the isolated U2AF2^Cy3/Cy5^ subunit.

**Figure 9.**
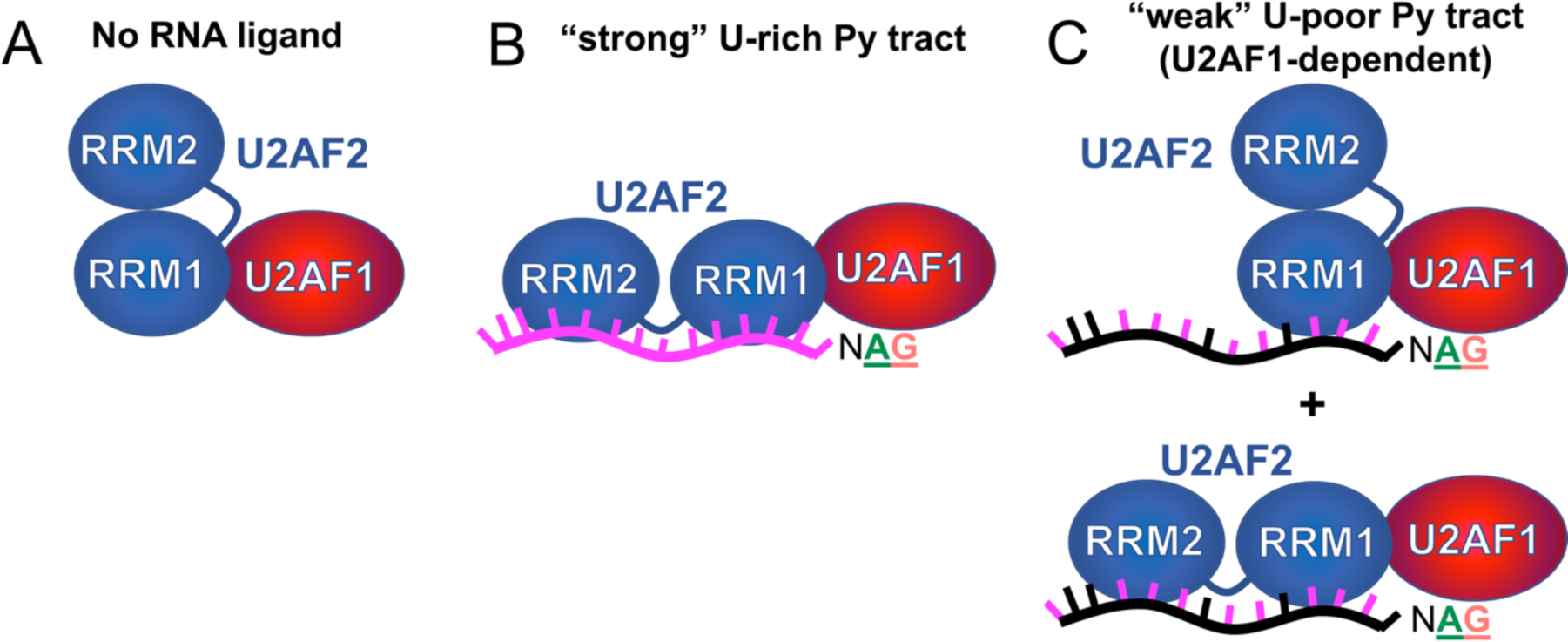
Models for splice site RNAs switching of the conformations of U2AF2 in the U2AF1– U2AF2 heterodimer. (**A**) The closed U2AF2 conformation predominates in the absence of RNA ligand. (**B**) Binding to a U-rich splice site RNA switches U2AF2 to the open, side-by-side RRM configurations. (**C**) A degenerate splice site signal with a short, interrupted Py tract preceding the AG-dinucleotide binds populations of closed and open conformations.

Out of the seven splice sites tested here, the S34F mutation induced large shifts in the populations of U2AF2 conformations for only two cases, *FMR1* and *IgM(−3U)*. The S34F mutation favored the open U2AF2 conformation when bound to the *FMR1* site but closed for the *IgM(−3U)* complex. These opposite shifts can be attributed to the context of the surrounding splice site sequences. For example, if the bulky phenylalanine side chain strains the closed U2AF2 conformation, the *FMR1* site includes an upstream U-rich track for docking of the open U2AF2 RRM2 whereas *IgM* lacks a uridine tract. Since the S34F mutation appears to block −3U/C sensing by the U2AF2 conformations, it is unsurprising that U2AF2 in the S34F mutant heterodimer primarily remains closed when bound to either *IgM* and *IgM(−3U)*, whereas the open U2AF2 population of the wild-type heterodimer increases following introduction of the −3U. We conclude that the S34F mutation can, but does not as a rule, influence the population of U2AF2 conformations. Alternatively, we suggest that S34F mutation alters the RNA binding characteristics of the U2AF1–U2AF2 heterodimer through direct RNA contacts, consistent with the location of this mutational hotspot at a putative RNA interface (20,26,27). Despite subtle effects of the S34F mutation on the apparent equilibrium RNA binding affinities of the human or fission yeast heterodimers (19,21), we show here that the S34F mutation dramatically increases the rates of fission yeast U2AF heterodimer interactions with *DEK* splice site variants. However, we caution that these RNA binding experiments with purified proteins are unlikely to directly correlate with splicing rates in cells, for which U2AF delivery and release from the splice site is governed by many factors including local concentrations, diffusion rates, and coupled gene expression events. Indeed, the S34F mutation of the human U2AF1 protein has been shown to delay splicing of representative transcripts until after release from the RNA polymerase complex (36).

A major outstanding question is whether one or both U2AF2 conformations are capable of promoting spliceosome assembly. One possibility is that only the open conformation of U2AF2 is active for spliceosome recruitment, whereas the closed conformation blocks splicing. Since the populations of each U2AF2 conformation depend on the degeneracy of the Py tract sequence, Py tract-sensing conformations with different activities could provide a checkpoint for splice site fidelity and also explain the different splicing efficiencies of uridine-rich and uridine-poor splice sites. The open U2AF2 population and hence splicing activity would be expected to increase following association of the U2AF heterodimer with uridine-rich Py tract signals such as *AdML.* Conversely if the closed fraction of U2AF2 molecules is inactive, a mixture of closed and open U2AF2 conformations would decrease the net splicing activity of uridine-poor Py tract signals. Although the U2AF2 conformations themselves do not appear tunable in response to RNA sequence (only the populations of two conformational classes), different ratios of active/inactive structures would enable U2AF2 to act as a graded rheostat for tunable splicing in response to different splice site signals.

Alternatively, it is possible that both open and closed U2AF2 conformations can initiate spliceosome assembly. Based on our detection of only two well-defined U2AF2 states among the RNA-free heterodimer and seven splice sites complexes, it appears unlikely that these conformations serve to adapt the U2AF2 structure to fit the broad range of human splice site signals. Yet, an intriguing possibility remains that the U2AF1-dependent class of uridine-poor splice sites (also called AG-dependent) may be spliced primarily through the closed U2AF2 conformation, whereas uridine-rich U2AF1-independent sites would be spliced *via* open U2AF2. In a minor U12-type spliceosome that entirely lacks Py tract signals, the U2AF1-like ZRSR2 subunit contains a heterodimerization domain that is expected to bind a U2AF2-like paralogue (9). One may suppose that a yet-to-be-identified U2AF2 subunit of the minor spliceosome would recognize the uridine-poor U12-type splice sites primarily in the closed conformation. Moreover, we suggest that the ratios of open/closed conformations observed here for human U2AF2 will differ from homologues that bind different consensus Py tracts. For example, we anticipate that the closed U2AF2 conformation will dominate recognition of the negligible or very short U-tracts of *Neurospora crassa* and *Caenorhabditis elegans* splice sites, which also are highly dependent on the U2AF1 subunit for binding and splicing (4,37,38).

In cells, a third splicing factor subunit called SF1 binds to U2AF2 as a ternary complex with U2AF1 and recognizes the branch site splicing signal upstream of the Py tract. The different geometries of the closed and open U2AF2 structures predict that the either proximal or distal branch sites would be chosen by SF1 depending on the population of these two U2AF2 conformations. If the uridine-poor, U2AF1-dependent splice sites rely on the closed U2AF2 conformation for splicing, then by extension, we expect the branch sites to be located closer to the spliced junctions of these sites. Accordingly, the ZRSR2-dependent branch sites of the U12 AT-AC introns typically are located closer to the spliced junction than those of the major, U2 GT-AG class in which the U2AF heterodimer functions (~12 *vs.* >20 nucleotides) (39). Moreover, the arginine-serine (RS)-rich effector domains of either U2AF1 or U2AF2 can promote U2 snRNA annealing with the branch site (10,40). We envision that the U2AF1 RS domain would drive U2 snRNA annealing with branch sites proximal to the U2AF1-dependent spliced junctions, raising the question of whether ZRSR2 supersedes the need for a U2AF2 paralogue for U12-type splicing. Here, we focused our studies on the minimal U2AF2 regions for U2AF1 heterodimerization and Py tract recognition, which avoids the potential complications of labeling full length U2AF2 in the presence of its the cysteine-rich SF1-interaction domain. It will be important to develop methods to study the conformational dynamics among the participating subunits of the intact U2AF1–U2AF2–SF1–splice site RNA complex, as well as to explore the conformational evolution of U2AF2 homologues and potential paralogues in the U12-type spliceosome.

## SUPPLEMENTARY DATA

Supplementary Data are available online.

### ACKNOWLEDGEMENTS

We are grateful to Prof. Stephanie Halene for discussions and sharing unpublished data.

### FUNDING

This work was supported by National Institutes of Health grants R01 GM070503 to C.L.K. and R01 GM099719 to D.N.E.

## REFERENCES

1. Kastner, B., Will, C.L., Stark, H. and Luhrmann, R. (2019) Structural insights into nuclear pre-mRNA splicing in higher eukaryotes. Cold Spring Harbor perspectives in biology.

2. Wu, S., Romfo, C.M., Nilsen, T.W. and Green, M.R. (1999) Functional recognition of the 3’ splice site AG by the splicing factor U2AF^35^. Nature, 402, 832–835.

3. Merendino, L., Guth, S., Bilbao, D., Martinez, C. and Valcarcel, J. (1999) Inhibition of msl-2 splicing by Sex-lethal reveals interaction between U2AF^35^ and the 3’ splice site AG. Nature, 402, 838–841.

4. Zorio, D.A. and Blumenthal, T. (1999) Both subunits of U2AF recognize the 3’ splice site in *Caenorhabditis elegans*. Nature, 402, 835–838.

5. Shao, C., Yang, B., Wu, T., Huang, J., Tang, P., Zhou, Y., Zhou, J., Qiu, J., Jiang, L., Li, H. et al. (2014) Mechanisms for U2AF to define 3’ splice sites and regulate alternative splicing in the human genome. Nat Struct Mol Biol, 21, 997–1005.

6. Fu, X.D. and Ares, M., Jr. (2014) Context-dependent control of alternative splicing by RNA-binding proteins. Nature reviews. Genetics, 15, 689–701.

7. Dvinge, H., Kim, E., Abdel-Wahab, O. and Bradley, R.K. (2016) RNA splicing factors as oncoproteins and tumour suppressors. Nat Rev Cancer, 16, 413–430.

8. Madan, V., Kanojia, D., Li, J., Okamoto, R., Sato-Otsubo, A., Kohlmann, A., Sanada, M., Grossmann, V., Sundaresan, J., Shiraishi, Y. et al. (2015) Aberrant splicing of U12-type introns is the hallmark of ZRSR2 mutant myelodysplastic syndrome. Nat Commun, 6, 6042.

9. Shen, H., Zheng, X., Luecke, S. and Green, M.R. (2010) The U2AF35-related protein Urp contacts the 3’ splice site to promote U12-type intron splicing and the second step of U2-type intron splicing. Genes Dev, 24, 2389–2394.

10. Shen, H. and Green, M.R. (2004) A pathway of sequential arginine-serine-rich domain-splicing signal interactions during mammalian spliceosome assembly. Mol Cell, 16, 363–373.

11. Shen, H., Kan, J.L. and Green, M.R. (2004) Arginine-serine-rich domains bound at splicing enhancers contact the branchpoint to promote prespliceosome assembly. Mol Cell, 13, 367–376.

12. Shen, H. and Green, M.R. (2006) RS domains contact splicing signals and promote splicing by a common mechanism in yeast through humans. Genes Dev, 20, 1755–1765.

13. Thickman, K.R., Swenson, M.C., Kabogo, J.M., Gryczynski, Z. and Kielkopf, C.L. (2006) Multiple U2AF^65^ binding sites within SF3b155: Thermodynamic and spectroscopic characterization of protein-protein interactions among pre-mRNA splicing factors. J Mol Biol, 356, 664–683.

14. Cass, D.M. and Berglund, J.A. (2006) The SF3b155 N-terminal domain is a scaffold important for splicing. Biochemistry, 45, 10092–10101.

15. Yoshida, K., Sanada, M., Shiraishi, Y., Nowak, D., Nagata, Y., Yamamoto, R., Sato, Y., Sato-Otsubo, A., Kon, A., Nagasaki, M. et al. (2011) Frequent pathway mutations of splicing machinery in myelodysplasia. Nature, 478, 64–69.

16. Graubert, T.A., Shen, D., Ding, L., Okeyo-Owuor, T., Lunn, C.L., Shao, J., Krysiak, K., Harris, C.C., Koboldt, D.C., Larson, D.E. et al. (2012) Recurrent mutations in the U2AF1 splicing factor in myelodysplastic syndromes. Nat Genet, 44, 53–57.

17. Imielinski, M., Berger, A.H., Hammerman, P.S., Hernandez, B., Pugh, T.J., Hodis, E., Cho, J., Suh, J., Capelletti, M., Sivachenko, A. et al. (2012) Mapping the hallmarks of lung adenocarcinoma with massively parallel sequencing. Cell, 150, 1107–1120.

18. Wu, S.J., Tang, J.L., Lin, C.T., Kuo, Y.Y., Li, L.Y., Tseng, M.H., Huang, C.F., Lai, Y.J., Lee, F.Y., Liu, M.C. et al. (2013) Clinical implications of U2AF1 mutation in patients with myelodysplastic syndrome and its stability during disease progression. Am J Hematol, 88, E277–282.

19. Okeyo-Owuor, T., White, B.S., Chatrikhi, R., Mohan, D.R., Kim, S., Griffith, M., Ding, L., Ketkar-Kulkarni, S., Hundal, J., Laird, K.M. et al. (2015) U2AF1 mutations alter sequence specificity of pre-mRNA binding and splicing. Leukemia, 29, 909–917.

20. Ilagan, J.O., Ramakrishnan, A., Hayes, B., Murphy, M.E., Zebari, A.S., Bradley, P. and Bradley, R.K. (2015) U2AF1 mutations alter splice site recognition in hematological malignancies. Genome Res, 25, 14–26.

21. Fei, D.L., Motowski, H., Chatrikhi, R., Prasad, S., Yu, J., Gao, S., Kielkopf, C.L., Bradley, R.K. and Varmus, H. (2016) Wild-type U2AF1 antagonizes the splicing program characteristic of U2AF1-mutant tumors and Is required for cell survival. PLoS Genet, 12, e1006384.

22. Glasser, E., Agrawal, A.A., Jenkins, J.L. and Kielkopf, C.L. (2017) Cancer-associated mutations mapped on high-resolution structures of the U2AF2 RNA recognition motifs. Biochemistry, 56, 4757–4761.

23. Yan, C., Wan, R. and Shi, Y. (2019) Molecular Mechanisms of pre-mRNA Splicing through Structural Biology of the Spliceosome. Cold Spring Harbor perspectives in biology, 11.

24. Loerch, S. and Kielkopf, C.L. (2016) Unmasking the U2AF homology motif family: a bona fide protein-protein interaction motif in disguise. RNA, 22, 1795–1807.

25. Kielkopf, C.L., Rodionova, N.A., Green, M.R. and Burley, S.K. (2001) A novel peptide recognition mode revealed by the X-ray structure of a core U2AF^35^/U2AF^65^ heterodimer. Cell, 106, 595–605.

26. Yoshida, H., Park, S.Y., Oda, T., Akiyoshi, T., Sato, M., Shirouzu, M., Tsuda, K., Kuwasako, K., Unzai, S., Muto, Y. et al. (2015) A novel 3’ splice site recognition by the two zinc fingers in the U2AF small subunit. Genes Dev, 29, 1649–1660.

27. Jenkins, J.L. and Kielkopf, C.L. (2017) Splicing factor mutations in myelodysplasias: Insights from spliceosome structures. Trends Genet, 33, 336–348.

28. Agrawal, A.A., Salsi, E., Chatrikhi, R., Henderson, S., Jenkins, J.L., Green, M.R., Ermolenko, D.N. and Kielkopf, C.L. (2016) An extended U2AF^65^-RNA-binding domain recognizes the 3’ splice site signal. Nat Commun, 7, 10950.

29. Mackereth, C.D., Madl, T., Bonnal, S., Simon, B., Zanier, K., Gasch, A., Rybin, V., Valcarcel, J. and Sattler, M. (2011) Multi-domain conformational selection underlies pre-mRNA splicing regulation by U2AF. Nature, 475, 408–411.

30. Jenkins, J.L., Agrawal, A.A., Gupta, A., Green, M.R. and Kielkopf, C.L. (2013) U2AF^65^ adapts to diverse pre-mRNA splice sites through conformational selection of specific and promiscuous RNA recognition motifs. Nucleic Acids Res, 41, 3859–3873.

31. Agrawal, A.A., McLaughlin, K.J., Jenkins, J.L. and Kielkopf, C.L. (2014) Structure-guided U2AF^65^ variant improves recognition and splicing of a defective pre-mRNA. Proc Natl Acad Sci U S A, 111, 17420–17425.

32. Jenkins, J.L., Laird, K.M. and Kielkopf, C.L. (2012) A broad range of conformations contribute to the solution ensemble of the essential splicing factor U2AF^65^. Biochemistry, 51, 5223–5225.

33. Voith von Voithenberg, L., Sanchez-Rico, C., Kang, H.S., Madl, T., Zanier, K., Barth, A., Warner, L.R., Sattler, M. and Lamb, D.C. (2016) Recognition of the 3’ splice site RNA by the U2AF heterodimer involves a dynamic population shift. Proc Natl Acad Sci U S A, 113, E7169–E7175.

34. Jenkins, J.L., Shen, H., Green, M.R. and Kielkopf, C.L. (2008) Solution conformation and thermodynamic characteristics of RNA binding by the splicing factor U2AF^65^. J Biol Chem, 283, 33641–33649.

35. Guth, S., Martinez, C., Gaur, R.K. and Valcarcel, J. (1999) Evidence for substrate-specific requirement of the splicing factor U2AF^35^ and for its function after polypyrimidine tract recognition by U2AF^65^. Mol Cell Biol, 19, 8263–8271.

36. Coulon, A., Ferguson, M.L., de Turris, V., Palangat, M., Chow, C.C. and Larson, D.R. (2014) Kinetic competition during the transcription cycle results in stochastic RNA processing. Elife, 3, e03939.

37. Henscheid, K.L., Voelker, R.B. and Berglund, J.A. (2008) Alternative modes of binding by U2AF^65^ at the polypyrimidine tract. Biochemistry, 47, 449–459.

38. Hollins, C., Zorio, D.A., MacMorris, M. and Blumenthal, T. (2005) U2AF binding selects for the high conservation of the C. elegans 3’ splice site. RNA, 11, 248–253.

39. Sharp, P.A. and Burge, C.B. (1997) Classification of introns: U2-type or U12-type. Cell, 91, 875–879.

40. Rudner, D.Z., Breger, K.S., Kanaar, R., Adams, M.D. and Rio, D.C. (1998) RNA binding activity of heterodimeric splicing factor U2AF: at least one RS domain is required for high-affinity binding. Mol Cell Biol, 18, 4004–4011.

