## Supplemental for "A Splice Site-Sensing Conformational Switch in U2AF2 is Modulated by U2AF1 and its Recurrent Myelodysplasia-Associated Mutation"

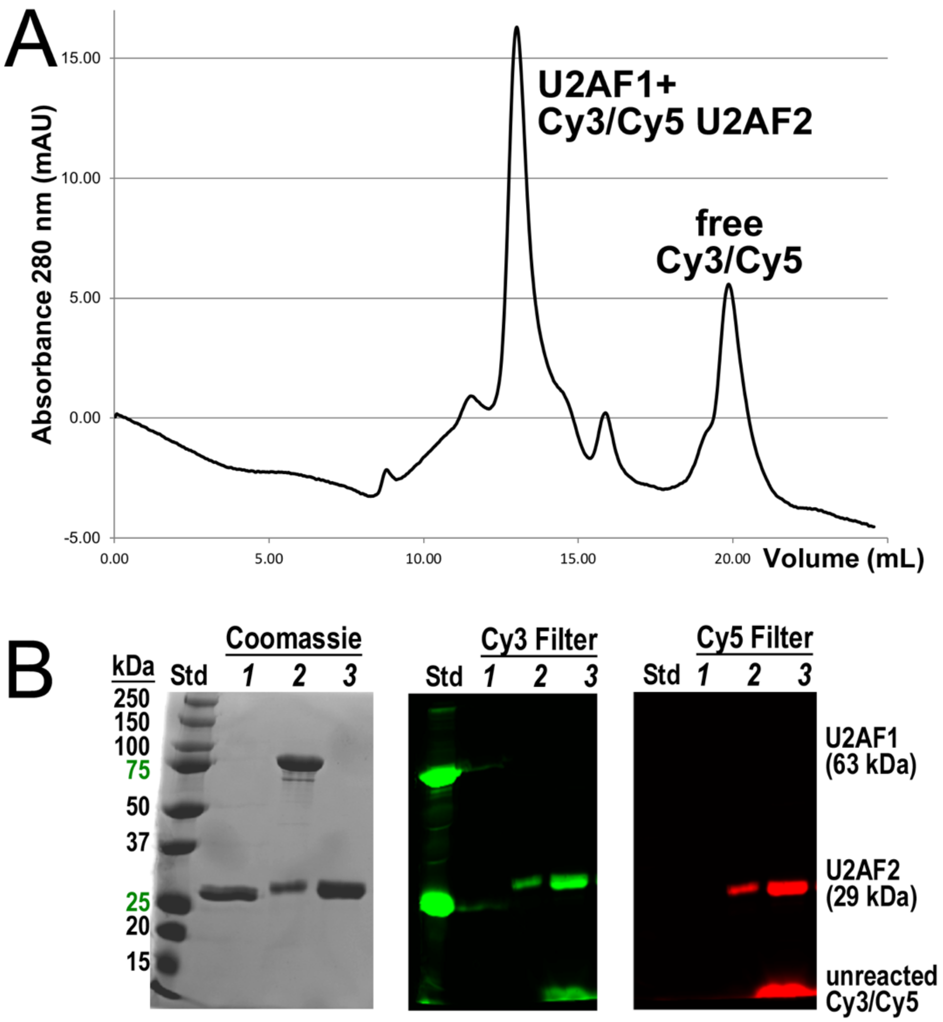


**Supplementary Figure S1.** Final preparation of 6xHisSumoMBP-U2AF1–U2AF2$^{Cy3/Cy5}$ heterodimer for smFRET. **(A)** Chromatogram of 6xHisSumoMBP-U2AF1 mixed with U2AF2$^{Cy3/Cy5}$ and separated from unreacted dye using a Superdex-200 Increase column. **(B)** SDS-PAGE of *1,* unlabeled U2AF2; *2,* final 6xHisSumoMBP-U2AF1–U2AF2$^{Cy3/Cy5}$ following size exclusion chromatography; *3*, unpurified U2AF2$^{Cy3/Cy5}$ labeling reaction, imaged with Coomassie blue stain, Cy3 filter, or Cy5 filter. A mixture of Cy3/Cy5 is expected at each U2AF2 site.

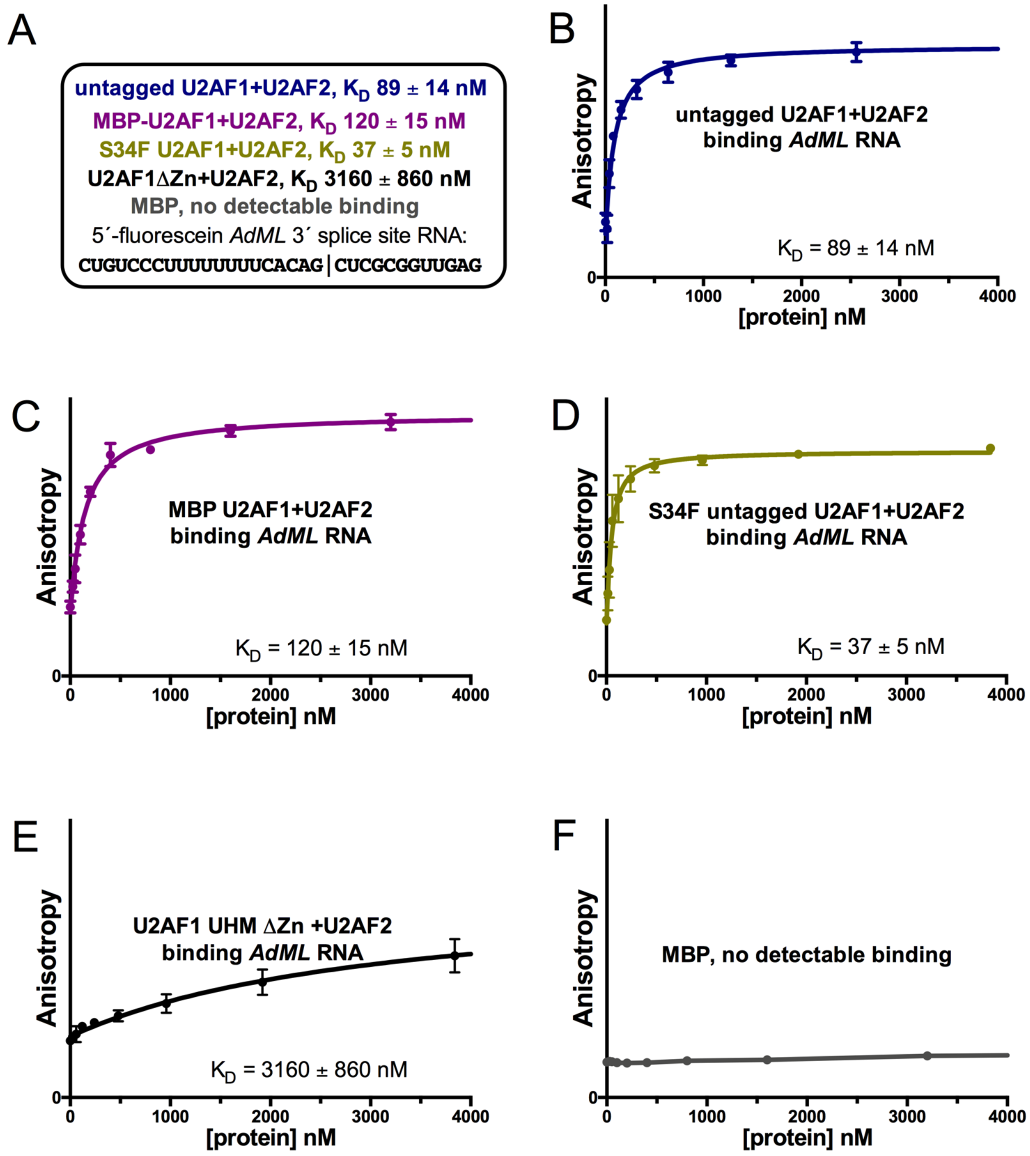


**Supplementary Figure S2.** Fluorescence anisotropy curves corresponding to the affinities shown in Figure 1F. (**A**) Apparent equilibrium dissociation constants ($K_D$) and standard deviations of three replicated *AdML* RNA binding experiments for each wild-type U2AF heterodimer variant and six for the S34F mutant, including (**B**) untagged heterodimer, (**C**) MBP-tagged U2AF1+U2AF2 heterodimer, (**D**) untagged S34F-mutant heterodimer, (**E**) untagged U2AF1 UHM domain (without zinc knuckles) heterodimer +U2AF2, or (**F**) the isolated MBP tag.

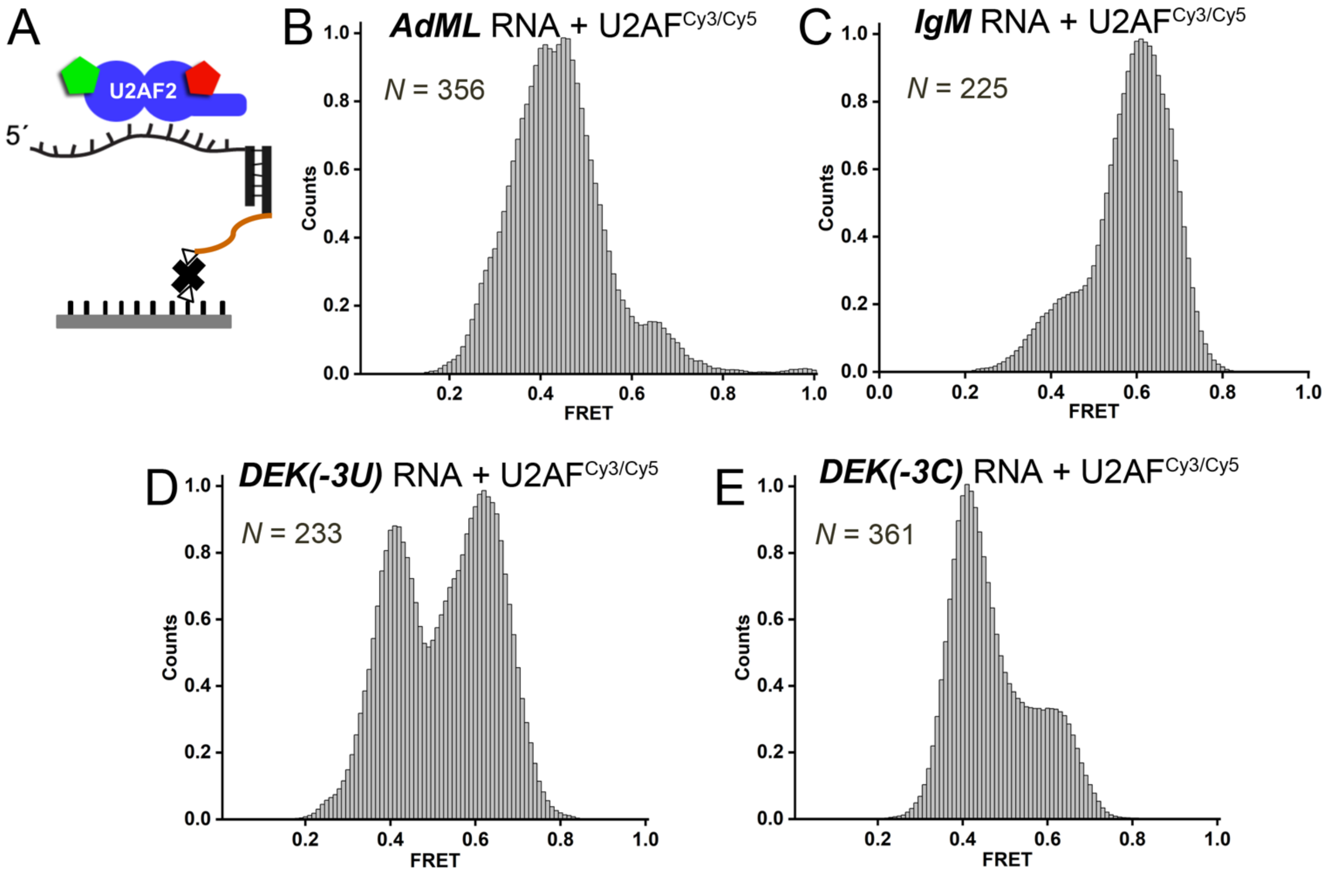


**Supplementary Figure S3.** smFRET of RNA complexes with the isolated U2AF2$^{Cy3/Cy5}$ subunit. (**A**) Scheme for splice site RNA immobilization and U2AF2$^{Cy3/Cy5}$ protein addition. (**B** - **E**) Histograms showing the distribution of FRET values for the untethered U2AF2$^{Cy3/Cy5}$ subunit bound to the indicated, slide-tethered RNA.

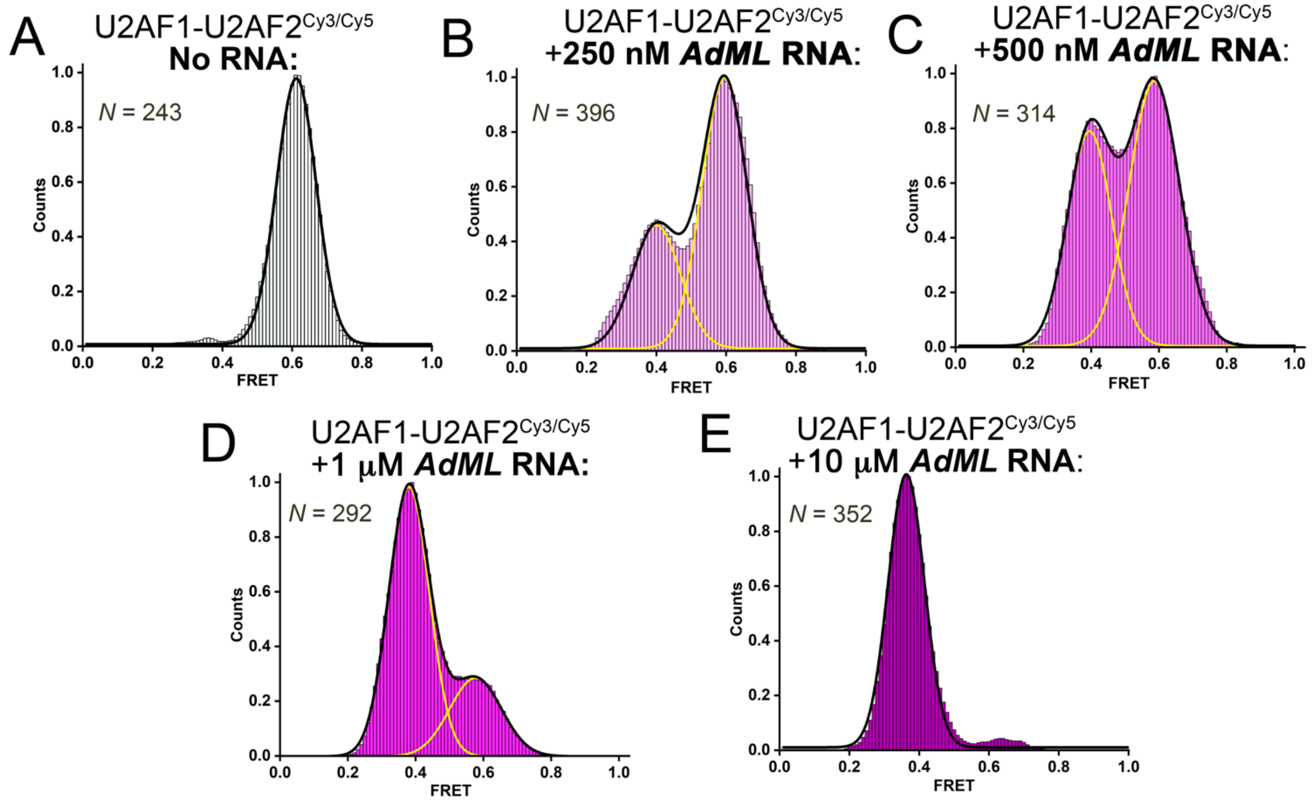


**Supplementary Figure S4.** Titration of immobilized U2AF1–U2AF2$^{Cy3/Cy5}$ heterodimer with the indicated amounts of *AdML* splice site RNA increases the population of U2AF2$^{Cy3/Cy5}$ to a lower FRET state. Black or yellow lines indicate the respective summed or individual Gaussian fits of the histograms.

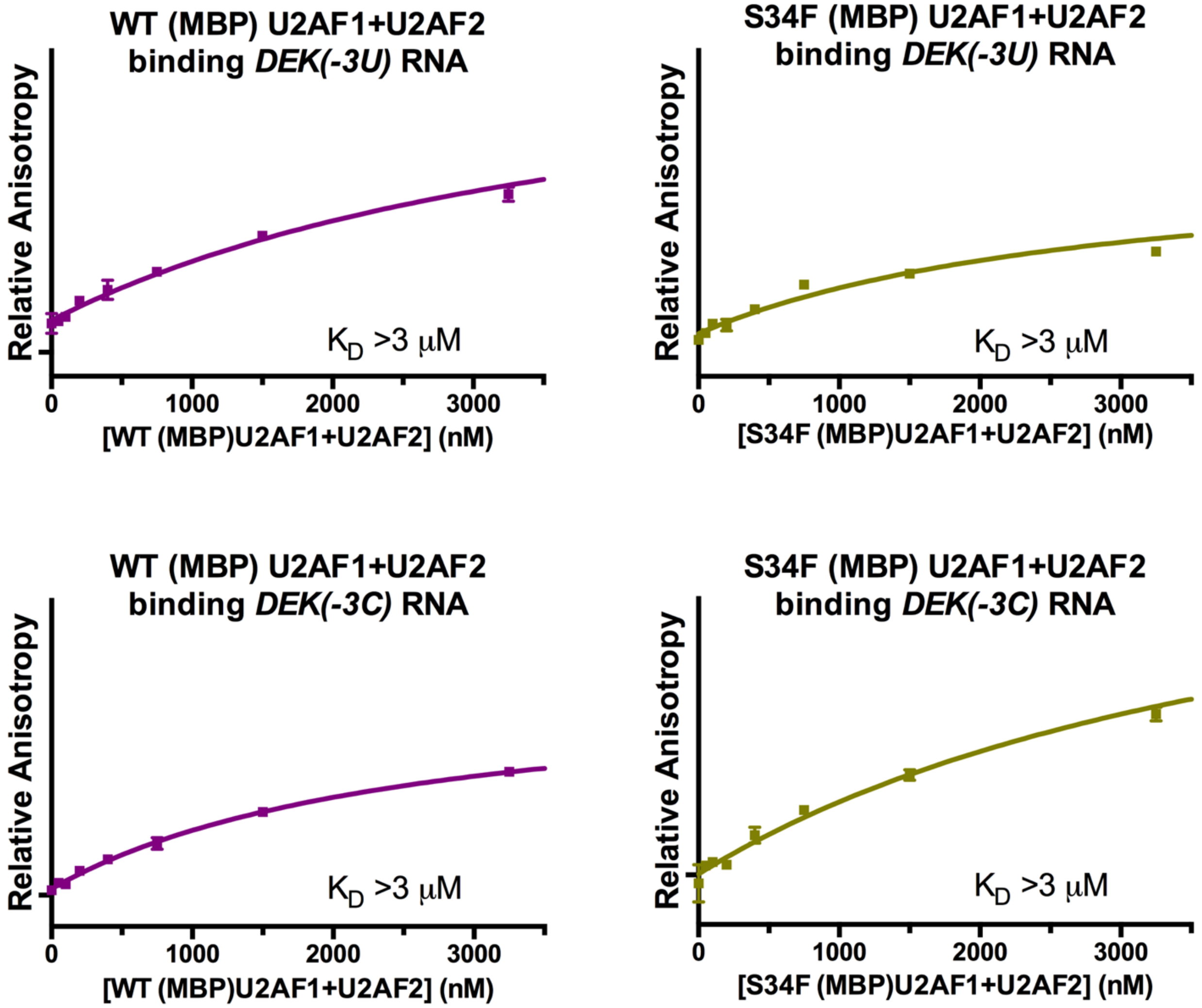


**Supplementary Figure S5.** Representative fluorescence anisotropy curves for human MBP-tagged U2AF1+U2AF2 heterodimer binding "weak" *DEK(-3U)* or *DEK(-3C)* splice site RNAs. The RNA oligonucleotide sequences are identical to those tested with the untagged SF1+U2AF1+U2AF2 ternary complex in {Okeyo-Owuor, 2015 #2361}. The apparent equilibrium dissociation constants ($K_D$) are beyond the limits of the accessible protein concentrations due to the very low binding affinities of the heterodimer for these non-consensus splice sites.

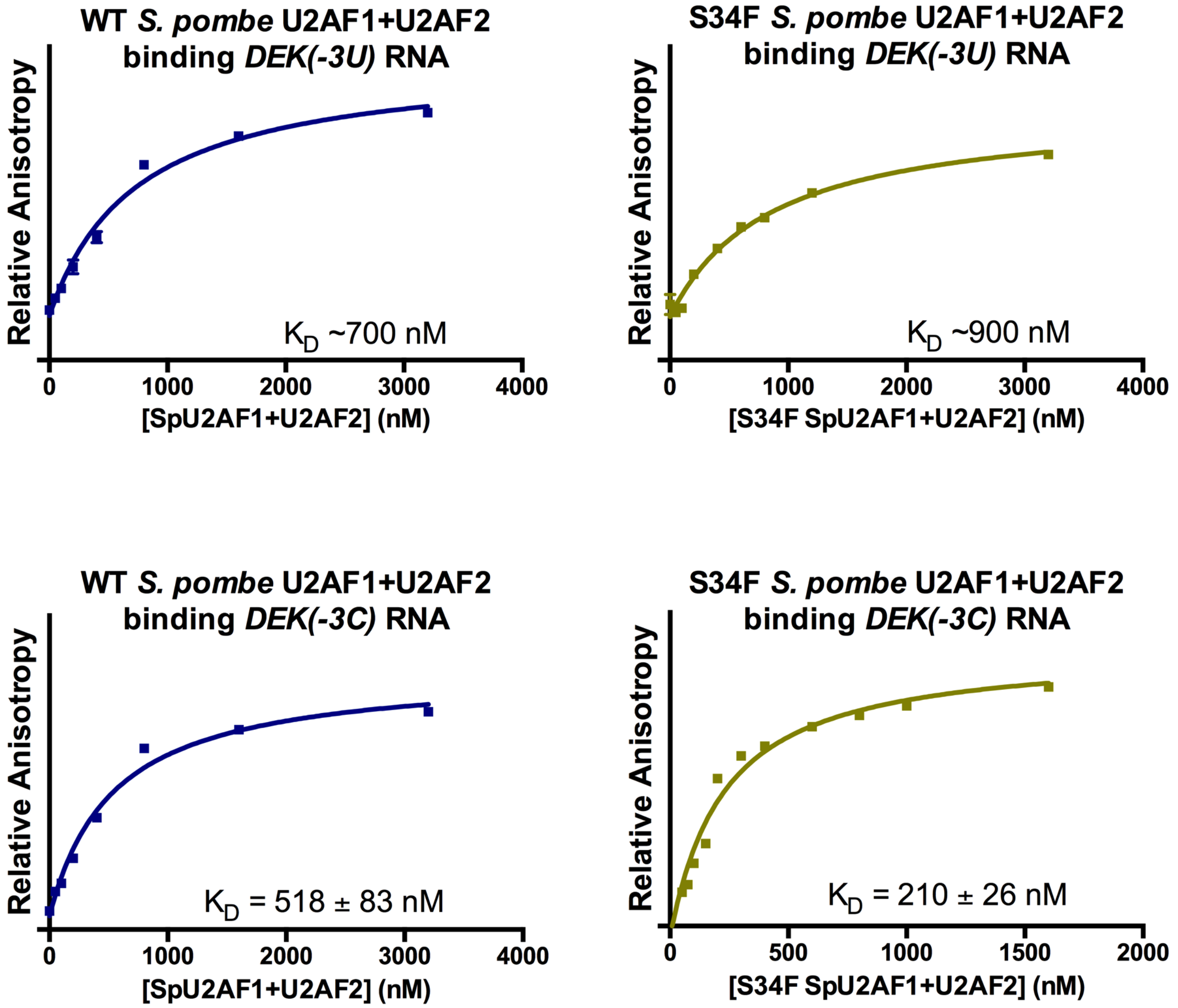


**Supplementary Figure S6.** Fluorescence anisotropy curves for wild-type (WT) or S34F-substituted *Schizosaccharomyces pombe* U2AF1+U2AF2 heterodimer binding *DEK(-3U/C)* splice site RNAs. The average data points and errors of three replicates are plotted. The apparent equilibrium dissociation constants ($K_D$) and errors (inset) are derived from two independent protein preparations of three technical triplicates each. The $K_D$ values for the *DEK* splice site are estimates due to the low binding affinity and impractical protein concentrations for curve saturation.
